# *IER5*, a DNA-damage response gene, is required for Notch-mediated induction of squamous cell differentiation

**DOI:** 10.1101/2020.04.21.051730

**Authors:** Li Pan, Madeleine E. Lemieux, Tom Thomas, Julia M. Rogers, Winston Lee, Carl Johnson, Lynette Sholl, Andrew P. South, Jarrod A. Marto, Guillaume O. Adelmant, Stephen C. Blacklow, Jon C. Aster

**Author notes:** Department of Pathology, University of Michigan Medical School, Ann Arbor, MI.

## Abstract

Notch signaling regulates normal squamous cell proliferation and differentiation and is frequently disrupted in squamous cell carcinomas, in which Notch is a key tumor suppressive pathway. To identify the direct targets of Notch that produce these phenotypes, we introduced a conditional Notch transgene into squamous cell carcinoma cell lines, which respond to Notch activation in 2D culture and in organoid cultures by undergoing differentiation. RNA-seq and ChIP-seq analyses show that in squamous cells Notch activates a context-specific program of gene expression that depends on lineage-specific regulatory elements, most of which lie in long-range enhancers. Among the direct Notch target genes are multiple DNA damage response genes, including *IER5*, which is regulated by Notch through several enhancer elements. We show that *IER5* is required for Notch-induced differentiation in squamous carcinoma cells and in TERT-immortalized keratinocytes. Its function is epistatic to *PPP2R2A*, which encodes the B55*α* subunit of PP2A, and IER5 interacts with B55*α* in cells and in purified systems. These results show that Notch and DNA-damage response pathways converge in squamous cells and that some components of these pathways promote differentiation, which may serve to eliminate DNA-damaged cells from the proliferative pool in squamous epithelia. Crosstalk involving Notch and PP2A may enable Notch signaling to be tuned and integrated with other pathways that regulate squamous differentiation. Our work also suggests that squamous cell carcinomas remain responsive to Notch signaling, providing a rationale for reactivation of Notch as a therapeutic strategy in these cancers.

**Impact:** Our findings highlight context-specific crosstalk between Notch, DNA damage response genes, and PP2A, and provide a roadmap for understanding how Notch induces the differentiation of squamous cells.

## Introduction

Notch receptors participate in a conserved signaling pathway in which successive ligand-mediated proteolytic cleavages by ADAM10 and *γ*-secretase permit intracellular Notch (ICN) to translocate to the nucleus and form a Notch transcription complex (NTC) with the DNA binding factor RBPJ and co-activators of the Mastermind-like (MAML) family (for review, see (Bray, 2016)). Outcomes of Notch activation are dose and cell context-dependent, in part because most Notch response elements lie within lineage-specific enhancers (Castel et al., 2013; Ryan et al., 2017; Skalska et al., 2015; H. Wang et al., 2014). As a result, Notch-dependent transcriptional programs vary widely across cell types.

The context-dependency of outcomes produced by Notch signaling is reflected in the varied patterns of Notch mutations that are found in different cancers (for review, see (Aster, Pear, & Blacklow, 2017)). In some cancers oncogenic gain-of-function Notch mutations predominate, but in human cutaneous squamous cell carcinoma (SCC) (South et al., 2014; N. J. Wang et al., 2011) loss-of-function mutations are common, early driver events, observations presaged by work showing that loss of Notch function promotes skin cancer development in mouse models (Nicolas et al., 2003; Proweller et al., 2006). The mechanism underlying the tumor suppressive effect of Notch appears to involve its ability to promote squamous differentiation at the expense of self-renewal, a function that is operative in other squamous epithelia (Alcolea et al., 2014), where Notch also has tumor suppressive activities (Agrawal et al., 2011; Agrawal et al., 2012; Loganathan et al., 2020). In line with this idea, conditional ablation of *Notch1* in postnatal mice results in epidermal hyperplasia and expansion of proliferating basal-like cells (Nicolas et al., 2003; Rangarajan et al., 2001). Moreover, murine and human *β*-papilloma viruses express E6 proteins that target MAML1 and inhibit Notch function (Meyers, Uberoi, Grace, Lambert, & Munger, 2017; Tan et al., 2012), thereby causing epidermal hyperplasia and delayed differentiation of infected keratinocytes. Conversely, constitutively active forms of Notch enhance keratinocyte differentiation *in vitro* and *in vivo* (Nickoloff et al., 2002; Rangarajan et al., 2001; Uyttendaele, Panteleyev, de Berker, Tobin, & Christiano, 2004).

While these studies delineate a pro-differentiation, tumor suppressive role for Notch in squamous cells, little is known about the Notch target genes that confer this phenotype. Work to date has focused on candidate genes chosen for their known activities in keratinocytes or their roles as Notch target genes in other cell types. These include *CDKN1A*/p21 (Rangarajan et al., 2001), which has been linked to cell cycle arrest and differentiation (Missero, Di Cunto, Kiyokawa, Koff, & Dotto, 1996); *HES1*, which represses basal fate/self-renewal (Blanpain, Lowry, Pasolli, & Fuchs, 2006); and *IRF6*, expression of which positively correlates with Notch activation in keratinocytes (Restivo et al., 2011). However, dose- and time-controlled genome-wide studies to determine the immediate, direct effects of Notch activation in squamous-lineage cells have yet to be performed.

To this end, we developed and validated 2D and 3D culture models of malignant and non-transformed human squamous epithelial cells in which regulated Notch activation produces growth arrest and squamous differentiation. We find that immediate, direct Notch target genes are largely keratinocyte-specific and are associated with lineage-specific NTC-binding enhancers enriched for the motifs of transcription factors linked to regulation of keratinocyte differentiation, particularly AP1. Among these genes are multiple genes previously shown to be upregulated by DNA damage and cell stress, including *IER5*, a member of the AP1-regulated immediate early response gene family (Williams et al., 1999). Here we show that *IER5* is required for Notch-induced differentiation of human SCC cells and TERT-immortalized human keratinocytes, and that this requirement is abolished by knockout of the B55*α* regulatory subunit of PP2A, to which IER5 directly binds. Our studies provide the first genome-wide view of the effects of Notch on gene expression in cutaneous squamous carcinoma cells, highlights previously unrecognized crosstalk between Notch and DNA response genes, and point to the existence of a Notch-IER5-PP2A signaling axis that coordinates keratinocyte differentiation.

### Establishment of a Conditional Notch-on SCC Model

Timed, regulated ligand-mediated Notch activation is difficult to achieve because soluble Notch ligands do not induce signaling (Sun & Artavanis-Tsakonas, 1997). To bypass this limitation, we previously developed a simple method that relies on *γ*-secretase inhibitor (GSI) washout (Figure 1A), which delivers ICN to the nuclei of cells expressing mutated or truncated forms of membrane-tethered Notch in 15-30 minutes (Petrovic et al., 2019; Ryan et al., 2017; H. Wang et al., 2014; Weng et al., 2006). To create a SCC model in which GSI washout activates NOTCH1, we engineered a cDNA encoding a mutated truncated form of NOTCH1, ΔEGF-L1596H, that cannot respond to ligand and that has a point substitution in its negative regulatory region that produces ligand-independent, *γ*-secretase-dependent Notch activation (Gordon et al., 2009; Malecki et al., 2006). ΔEGF-L1596H was transduced into two human SCC cell lines, IC8 and SCCT2, that have biallelic inactivating mutations in *NOTCH1* and *TP53* (Inman et al., 2018), lesions that were confirmed by resequencing on a clinical-grade targeted exome NGS platform (summarized in Tables 1 and 2). In pilot studies, we observed that the growth of IC8 and SCCT2 cells transduced with empty virus was unaffected by the presence or absence of GSI, whereas the growth of lines transduced with ΔEGF-L1596H was reduced by GSI washout (Figure 1, Supplemental Figure 1A, B). As anticipated, GSI washout was accompanied by rapid activation of NOTCH1 (ICN1), which was followed by delayed upregulation of markers of differentiation, such as involucrin (Figure 1, Supplemental Figure 1C, D).

**Table 1.**
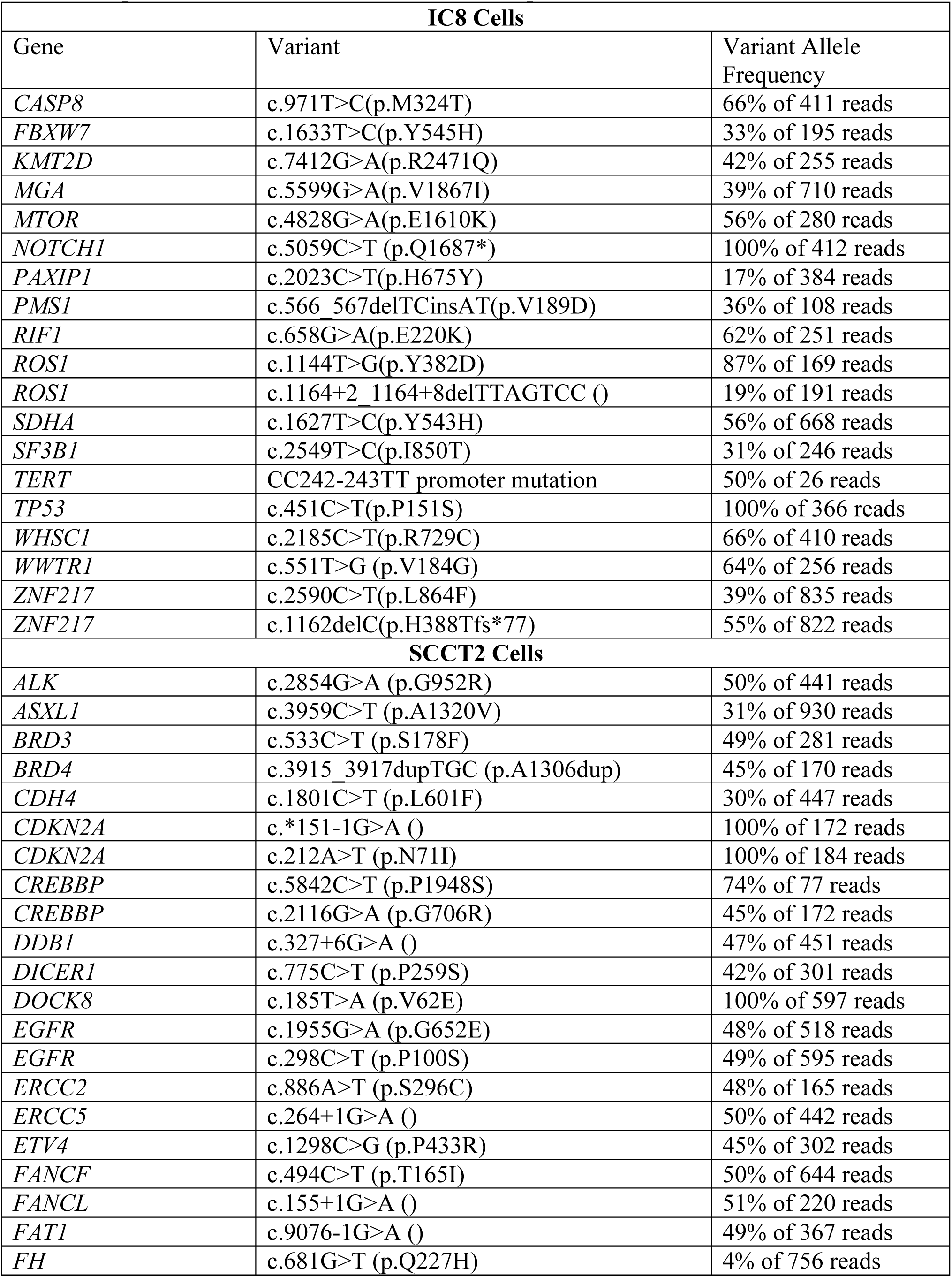
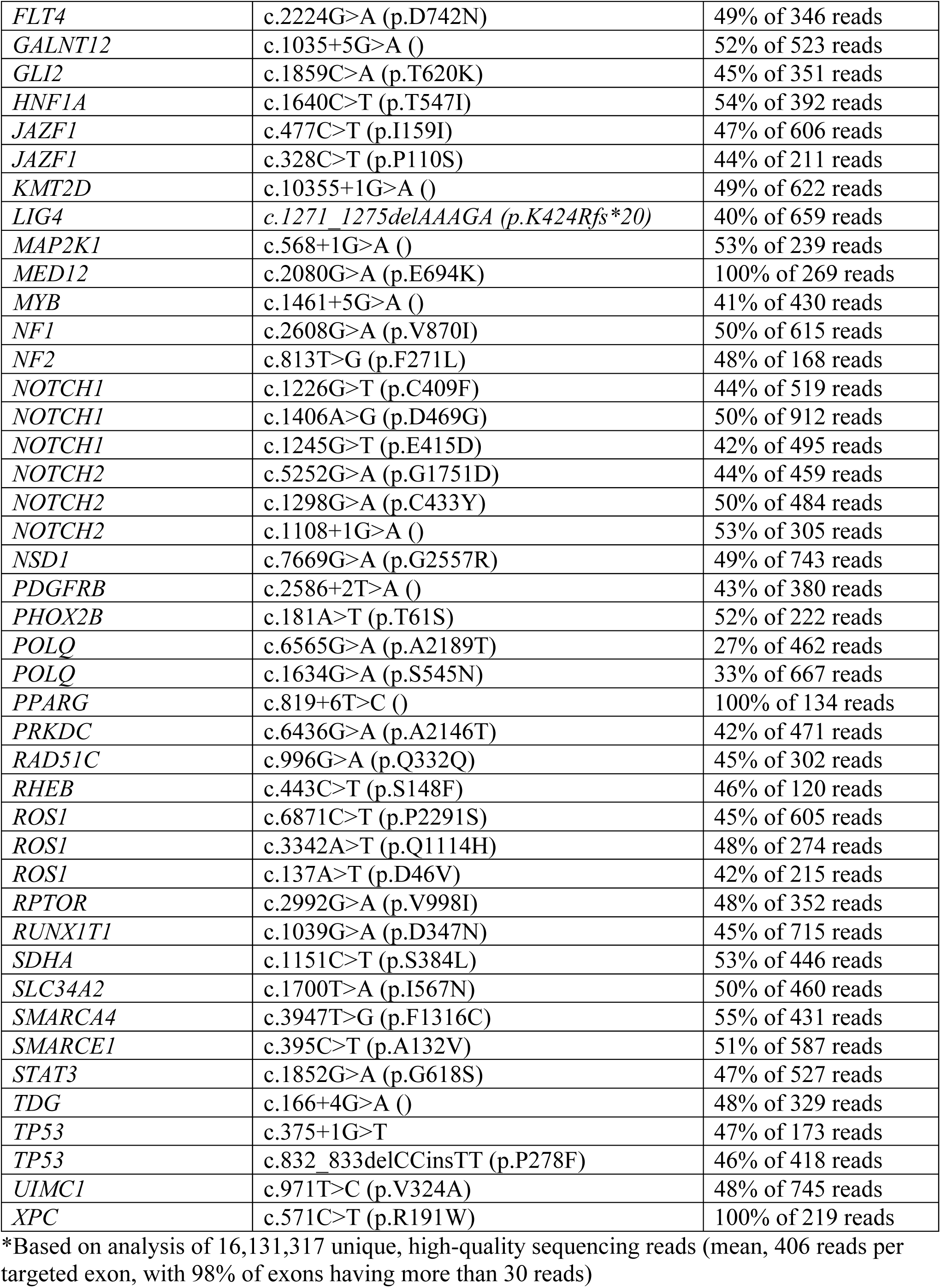
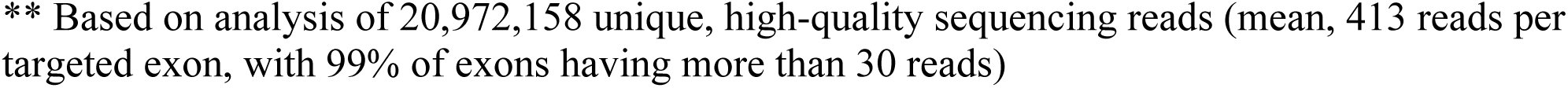
Sequence Variants, IC8* and SCCT2** Squamous Cell Carcinoma Cell Lines.

| IC8 Cells |  |  |
| --- | --- | --- |
| Gene | Variant | Variant Allele Frequency |
| *CASP8* | c.971T>C(p.M324T) | 66% of 411 reads |
| *FBXW7* | c.1633T>C(p.Y545H) | 33% of 195 reads |
| *KMT2D* | c.7412G>A(p.R2471Q) | 42% of 255 reads |
| *MGA* | c.5599G>A(p.V1867I) | 39% of 710 reads |
| *MTOR* | c.4828G>A(p.E1610K) | 56% of 280 reads |
| *NOTCH1* | c.5059C>T (p.Q1687*) | 100% of 412 reads |
| *PAXIP1* | c.2023C>T(p.H675Y) | 17% of 384 reads |
| *PMS1* | c.566_567delTCinsAT(p.V189D) | 36% of 108 reads |
| *RIF1* | c.658G>A(p.E220K) | 62% of 251 reads |
| *ROS1* | c.1144T>G(p.Y382D) | 87% of 169 reads |
| *ROS1* | c.1164+2_1164+8delTTAGTCC () | 19% of 191 reads |
| *SDHA* | c.1627T>C(p.Y543H) | 56% of 668 reads |
| *SF3B1* | c.2549T>C(p.I850T) | 31% of 246 reads |
| *TERT* | CC242-243TT promoter mutation | 50% of 26 reads |
| *TP53* | c.451C>T(p.P151S) | 100% of 366 reads |
| *WHSC1* | c.2185C>T(p.R729C) | 66% of 410 reads |
| *WWTR1* | c.551T>G (p.V184G) | 64% of 256 reads |
| *ZNF217* | c.2590C>T(p.L864F) | 39% of 835 reads |
| *ZNF217* | c.1162delC(p.H388Tfs*77) | 55% of 822 reads |
| SCCT2 Cells |  |  |
| *ALK* | c.2854G>A (p.G952R) | 50% of 441 reads |
| *ASXL1* | c.3959C>T (p.A1320V) | 31% of 930 reads |
| *BRD3* | c.533C>T (p.S178F) | 49% of 281 reads |
| *BRD4* | c.3915_3917dupTGC (p.A1306dup) | 45% of 170 reads |
| *CDH4* | c.1801C>T (p.L601F) | 30% of 447 reads |
| *CDKN2A* | c.*151-1G>A () | 100% of 172 reads |
| *CDKN2A* | c.212A>T (p.N71I) | 100% of 184 reads |
| *CREBBP* | c.5842C>T (p.P1948S) | 74% of 77 reads |
| *CREBBP* | c.2116G>A (p.G706R) | 45% of 172 reads |
| *DDB1* | c.327+6G>A () | 47% of 451 reads |
| *DICER1* | c.775C>T (p.P259S) | 42% of 301 reads |
| *DOCK8* | c.185T>A (p.V62E) | 100% of 597 reads |
| *EGFR* | c.1955G>A (p.G652E) | 48% of 518 reads |
| *EGFR* | c.298C>T (p.P100S) | 49% of 595 reads |
| *ERCC2* | c.886A>T (p.S296C) | 48% of 165 reads |
| *ERCC5* | c.264+1G>A () | 50% of 442 reads |
| *ETV4* | c.1298C>G (p.P433R) | 45% of 302 reads |
| *FANCF* | c.494C>T (p.T165I) | 50% of 644 reads |
| *FANCL* | c.155+1G>A () | 51% of 220 reads |
| *FAT1* | c.9076-1G>A () | 49% of 367 reads |
| *FH* | c.681G>T (p.Q227H) | 4% of 756 reads |
| *FLT4* | c.2224G>A (p.D742N) | 49% of 346 reads |
| *GALNT12* | c.1035+5G>A ( ) | 52% of 523 reads |
| *GLI2* | c.1859C>A (p.T620K) | 45% of 351 reads |
| *HNF1A* | c.1640C>T (p.T547I) | 54% of 392 reads |
| *JAZF1* | c.477C>T (p.I159I) | 47% of 606 reads |
| *JAZF1* | c.328C>T (p.P110S) | 44% of 211 reads |
| *KMT2D* | c.10355+1G>A ( ) | 49% of 622 reads |
| *LIG4* | c.1271_1275delAAAGA (p.K424Rfs*20) | 40% of 659 reads |
| *MAP2K1* | c.568+1G>A ( ) | 53% of 239 reads |
| *MED12* | c.2080G>A (p.E694K) | 100% of 269 reads |
| *MYB* | c.1461+5G>A ( ) | 41% of 430 reads |
| *NF1* | c.2608G>A (p.V870I) | 50% of 615 reads |
| *NF2* | c.813T>G (p.F271L) | 48% of 168 reads |
| *NOTCH1* | c.1226G>T (p.C409F) | 44% of 519 reads |
| *NOTCH1* | c.1406A>G (p.D469G) | 50% of 912 reads |
| *NOTCH1* | c.1245G>T (p.E415D) | 42% of 495 reads |
| *NOTCH2* | c.5252G>A (p.G1751D) | 44% of 459 reads |
| *NOTCH2* | c.1298G>A (p.C433Y) | 50% of 484 reads |
| *NOTCH2* | c.1108+1G>A ( ) | 53% of 305 reads |
| *NSD1* | c.7669G>A (p.G2557R) | 49% of 743 reads |
| *PDGFRB* | c.2586+2T>A ( ) | 43% of 380 reads |
| *PHOX2B* | c.181A>T (p.T61S) | 52% of 222 reads |
| *POLQ* | c.6565G>A (p.A2189T) | 27% of 462 reads |
| *POLQ* | c.1634G>A (p.S545N) | 33% of 667 reads |
| *PPARG* | c.819+6T>C ( ) | 100% of 134 reads |
| *PRKDC* | c.6436G>A (p.A2146T) | 42% of 471 reads |
| *RAD51C* | c.996G>A (p.Q332Q) | 45% of 302 reads |
| *RHEB* | c.443C>T (p.S148F) | 46% of 120 reads |
| *ROS1* | c.6871C>T (p.P2291S) | 45% of 605 reads |
| *ROS1* | c.3342A>T (p.Q1114H) | 48% of 274 reads |
| *ROS1* | c.137A>T (p.D46V) | 42% of 215 reads |
| *RPTOR* | c.2992G>A (p.V998I) | 48% of 352 reads |
| *RUNX1T1* | c.1039G>A (p.D347N) | 45% of 715 reads |
| *SDHA* | c.1151C>T (p.S384L) | 53% of 446 reads |
| *SLC34A2* | c.1700T>A (p.I567N) | 50% of 460 reads |
| *SMARCA4* | c.3947T>G (p.F1316C) | 55% of 431 reads |
| *SMARCE1* | c.395C>T (p.A132V) | 51% of 587 reads |
| *STAT3* | c.1852G>A (p.G618S) | 47% of 527 reads |
| *TDG* | c.166+4G>A ( ) | 48% of 329 reads |
| *TP53* | c.375+1G>T | 47% of 173 reads |
| *TP53* | c.832_833delCCinsTT (p.P278F) | 46% of 418 reads |
| *UIMC1* | c.971T>C (p.V324A) | 48% of 745 reads |
| *XPC* | c.571C>T (p.R191W) | 100% of 219 reads |

\*Based on analysis of 16,131,317 unique, high-quality sequencing reads (mean, 406 reads per targeted exon, with 98% of exons having more than 30 reads)
\*\* Based on analysis of 20,972,158 unique, high-quality sequencing reads (mean, 413 reads per targeted exon, with 99% of exons having more than 30 reads)

**Table 2.**
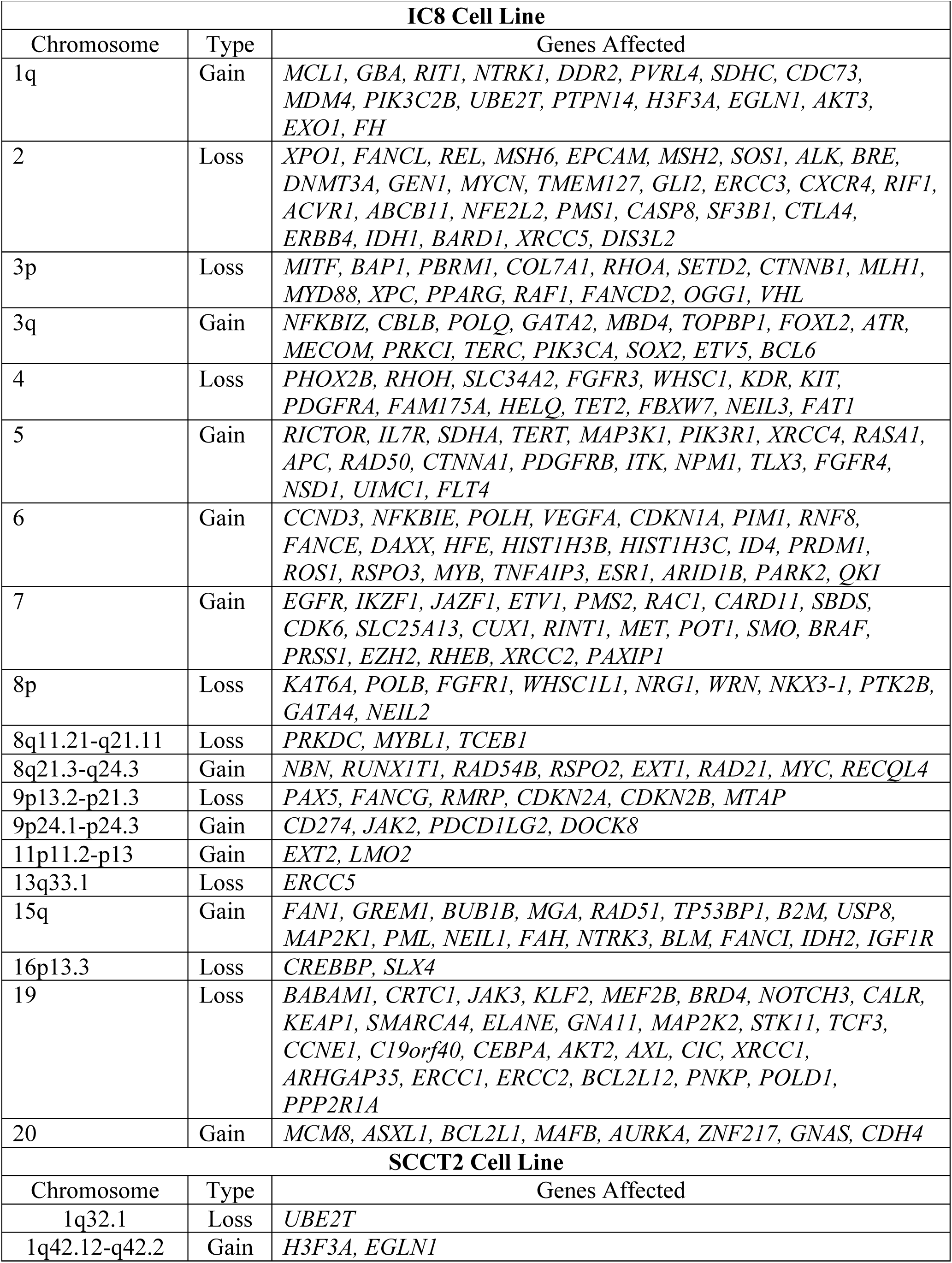
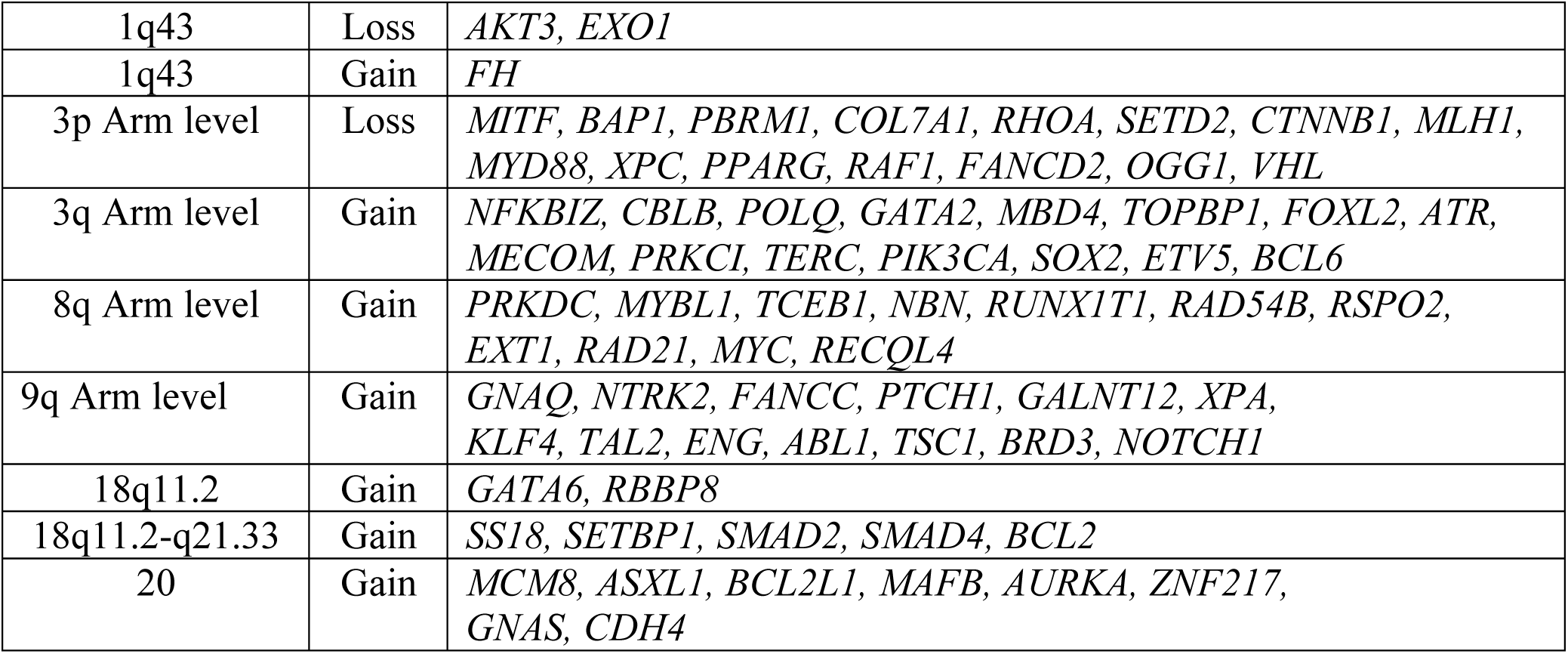
Copy Number Variants, Squamous Cell Carcinoma Cell Lines.

**Figure 1.**
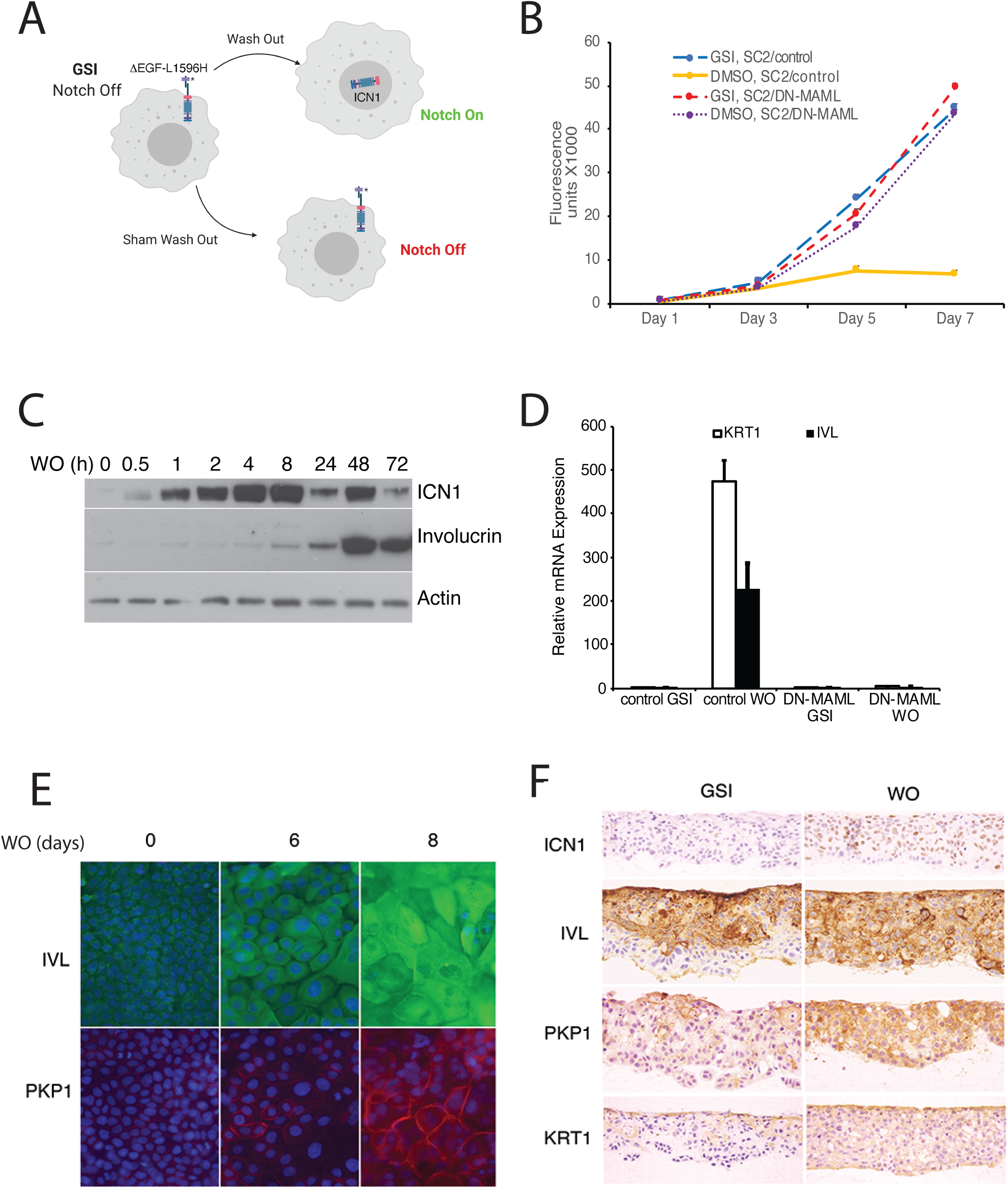
Notch activation induces growth arrest and differentiation of squamous carcinoma cells. (A) Strategy used to activate Notch in a regulated fashion. (B) Notch-induced suppression of SC2 cell growth in standard cultures is abrogated by DN-MAML, a specific inhibitor of canonical Notch signaling. SC2 cells were transduced with empty MigRI virus (con) or with MigRI virus encoding DN-MAML. Cell numbers at various times post-GSI washout (DMSO vehicle alone) or mock GSI-washout (GSI) were assessed using Cell Titer-Blue on independent cultures in quadruplicate. Error bars represent standard deviations. (C) Western blot showing the kinetics of ICN1 generation and increases in involucrin (IVL) following GSI washout in SC2 cells in standard cultures. (D) Notch-induced differentiation of SC2 cells is abrogated by DN- MAML. Transcripts for keratin1 (KRT1) and involucrin (IVL) were measured in the presence of GSI and 3 days after GSI washout (WO) in SC2 cells transduced with empty virus (con) or with DN-MAML. Transcript abundance was measured in experimental triplicates by RT-PCR and normalized against GAPDH. Error bars represent standard deviations of the mean. (E) Indirect immunofluorescence microscopy showing staining for involucrin (IVL, green) and plakophilin-1 (PKP1, red) in SC2 cells in 2D cultures at time 0 and 6 and 8 days after GSI washout (WO). Nuclei in each image were counterstained with DAPI. (F) Immunohistochemical staining of SC2 cells grown in raft cultures for 14 days in the presence of GSI or following GSI washout (WO). ICN1, activated intracellular NOTCH1.

To further characterize and optimize our system, we performed single cell-cloning of IC8*−*ΔEGF-L1596H cells and observed that growth arrest following GSI washout correlated with ICN1 accumulation (Figure 1, Supplemental Figure 2A, B). The subclone SC2, which showed moderate accumulation of ICN1 and sharply reduced growth following GSI washout, was selected for further study. We also observed that SC2 cells formed “skin-like” epithelia when seeded onto organotypic 3D cultures (Figure 1, Supplemental Figure 2C), whereas SCCT2- ΔEGF-L1596H cells did not (not shown); therefore, additional studies focused on IC8 cells and derivatives thereof. The ability of SC2 cells to form a multilayered epithelium in organotypic cultures in the Notch-on, growth suppressive state appears to stem from an emergent property, the Notch-dependent organization of these cells into a proliferating, ICN1-low basal layer in contact with matrix and a non-proliferating, ICN1-high suprabasal layer (Figure 1, Supplemental Figure 2C). Notably, growth arrest induced by GSI washout in SC2 cells was blocked by dominant-negative MAML1 (DN-MAML), a specific inhibitor of Notch-dependent transcription (Figure 1B) (Nam, Sliz, Pear, Aster, & Blacklow, 2007; Weng et al., 2003), and was accompanied by increases in multiple markers of squamous differentiation, such as involucrin, keratin1, and plakophilin1, in both 2D (Figure 1C-E) and in 3D cultures (Figure 1F).

**Figure 2.**
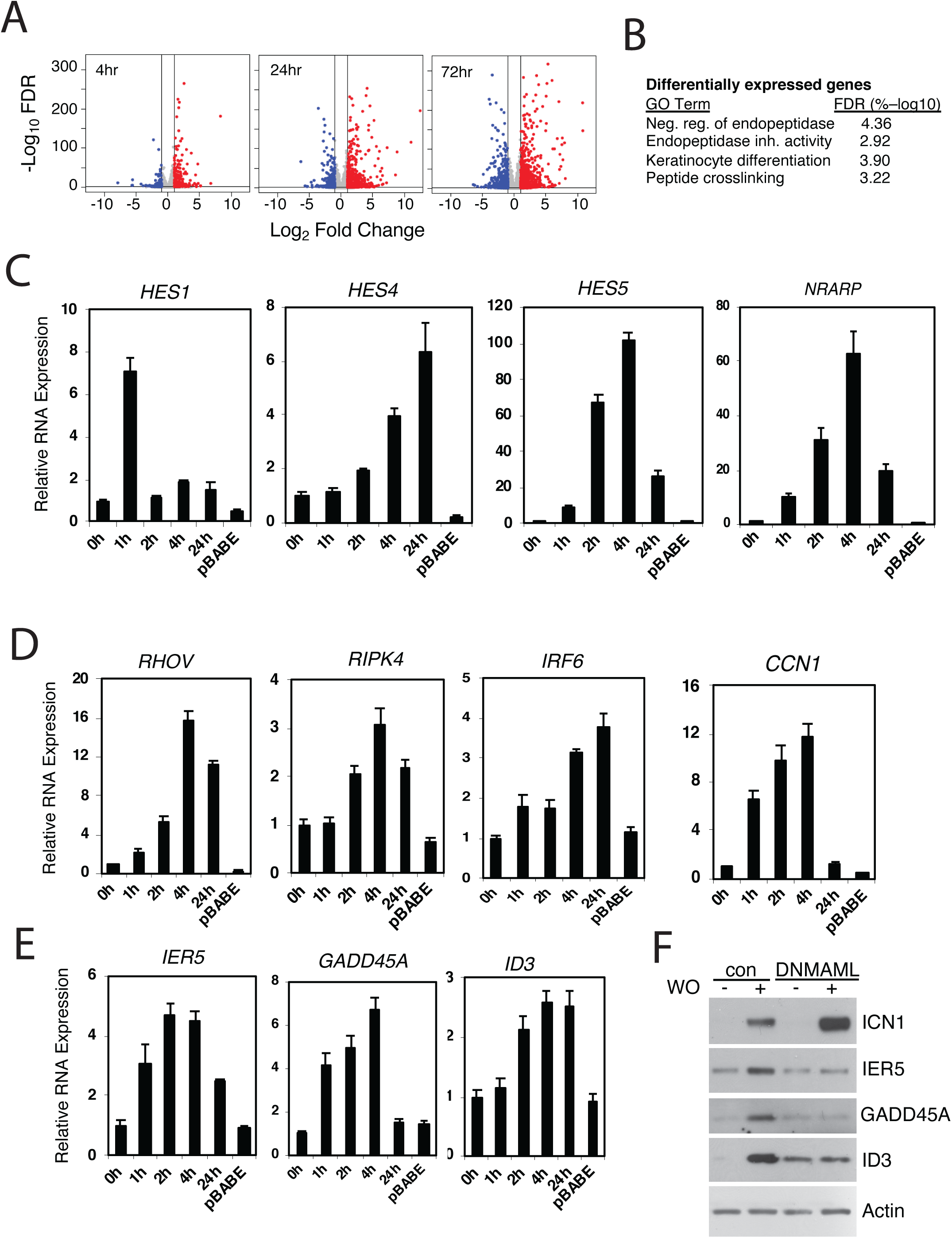
Identification of Notch-induced genes in squamous carcinoma cells. (A) Volcano plots showing changes in RNA transcript read counts induced by Notch activation in SC2 cells for 4, 24, and 72 h as compared to control cells treated with sham GSI washout. RNA-seq for each treatment group was performed in triplicate on independent cell cultures. Vertical lines denote a 2-fold change in read count, while the horizontal line denotes a false discovery rate (FDR) of 5%. (B) Clustered gene ontogeny (GO) annotation of differentially expressed genes in “Notch-on” SC2 cells. The most highly associated clustered GO terms are shown; other significant associated unclustered and clustered annotated gene sets (FDR<5%) are listed in Tables S4 and S5, respectively. (C-E) Transcriptional responses of selected “canonical” Notch target genes (C), genes linked to keratinocyte differentiation (D), and genes associated with DNA damage responses (E) to Notch activation in IC8-ΔEGF-L1596H cells. Transcript abundance was measure in experimental triplicates by RT-PCR and normalized against GAPDH. Error bars represent standard deviations of the mean. (F) Western blots of cell lysates prepared from IC8- ΔEGF-L1596H cells transduced with empty virus (con) or DN-MAML following sham GSI washout (-) or 24h post-GSI washout (+).

### Identification of a Notch-induced program of gene expression in SCC

To determine early and delayed effects of Notch on gene expression, we performed RNA-seq on SC2 cells in 2D cultures in the Notch-off state and following Notch activation (Figure 2A). Because *γ*-secretase has numerous substrates, as a control we performed RNA-seq on parental, non-transduced IC8 cells in the presence of GSI and following GSI washout, which revealed no significant GSI-dependent changes in gene expression in the absence of a Notch transgene, with significant being defined as adjusted p-value <0.0001 and log_2_ fold change >1 (Figure 2, Supplemental Figure 1). By contrast, Notch activation in SC2 cells produced significant changes in gene expression by 4h that became more pronounced by 24h and 72h (Figure 2A, see Tables S1-S3 for differentially expressed genes). Gene ontology (GO) analysis revealed enrichment among upregulated genes for those that are associated with keratinocyte differentiation and biology (Tables S4 and S5, summarized in Figure 2B). Among the rapidly upregulated genes (genes that increase in expression by 4h) were several reported targets of Notch in keratinocytes (e.g., *RHOV* (Pomrey & Radtke, 2010), *HES1* (Blanpain et al., 2006), and *IRF6* (Restivo et al., 2011)), but most genes in this class were novel and included: 1) direct targets of Notch in other lineages (e.g., *NRARP, HES4*, and *HES5*); 2) genes linked to keratinocyte differentiation (e.g., *RIPK4* (Kwa et al., 2014) and *SMAD3* (Meyers et al., 2017)); 3) genes associated with DNA damage responses in keratinocytes and other cell types (e.g., *GADD45A, CXCL8, IL1B, ID3, CYR61, BTG2, IER3*, and *IER5* (Kis et al., 2006; Kumar et al., 1998; Maeda et al., 2002; Rouault et al., 1996; Sesto, Navarro, Burslem, & Jorcano, 2002; Simbulan-Rosenthal et al., 2006)); and 4) genes associated with growth arrest of keratinocytes and other cell types (e.g., *HES1* (Blanpain et al., 2006), *GADD45A* (Maeda et al., 2002), and *BTG2* (Rouault et al., 1996)).

To confirm that these changes in gene expression are general features of Notch activation in IC8 cells (and not merely of the SC2 subclone) and to determine the kinetics of response, we performed RT-PCR analyses on a number of known and novel targets in pooled IC8-ΔEGF- L1596H transductants. This confirmed the Notch responsiveness of all genes tested, and also revealed variation in the kinetics of response, even among “canonical” Notch target genes. For example, *HES1* showed fast induction followed by rapid down-regulation, consistent with autoinhibition (Hirata et al., 2002), whereas *HES4, HES5*, and *NRARP* (a feedback inhibitor of NTC function (Jarrett et al., 2019)) showed more sustained increases in expression (Figure 2C). Genes encoding non-structural proteins known to be linked to squamous differentiation also were “early” responders (Figure 2D), as were genes linked to DNA damage/cell stress response (Figure 2E). In the case of the latter novel rapidly responding genes, we used DN-MAML blockade to confirm that protein levels also rose in a Notch-dependent fashion (Figure 2F). By contrast, increased expression of genes encoding structural proteins associated with keratinocyte differentiation (e.g., *IVL, KRT1, KRT13, KRT14, KRT16, KRT17, LOR, FLG*) was delayed, only emerging at 24-72h (Tables S1-S3). These findings suggest that Notch activation induces the expression of a core group of early direct target genes, setting in motion downstream events that lead to differentiation.

Notch activation also produced decreased expression of a smaller set of genes (Figure 2A, summarized in Tables S1-S3), possibly via induction of transcriptional repressors of the Hes family. These include multiple genes expressed by basal epidermal stem cells, including genes encoding the Notch ligand DLL1 (Lowell, Jones, Le Roux, Dunne, & Watt, 2000); *β*1-integrin (Jones & Watt, 1993); LRIG1 (Jensen & Watt, 2006), a negative regulator of epidermal growth factor receptor signaling; and multiple WNT ligands (WNT7A, 7B, 9A, 10A, and 11), of interest because WNT signaling contributes to maintenance of epidermal stem cells (Lim et al., 2013).

### Notch target genes are associated with lineage-specific NTC-binding enhancer elements

To identify sites of NTC-binding to Notch-responsive regulatory elements in IC8-ΔEGF-L1596H cells, we performed ChIP-seq for endogenous Notch transcription complex components (RBPJ and MAML1) 4h after Notch activation, as well as for RBPJ prior to Notch activation. In the Notch-on state, most MAML1 binding sites also bound RBPJ (8533/9,187 sites, 93%; Figure 3A), in line with studies showing that MAML1 association with DNA requires both RBPJ and NICD (Nam, Sliz, Song, Aster, & Blacklow, 2006). Approximately 92% of RBPJ/MAML1 co-binding sites (hereafter designated NTC binding sites) are in intergenic or intronic regions consistent with enhancers (Figure 3B). As predicted by past studies (Castel et al., 2013; Krejci & Bray, 2007; Ryan et al., 2017; H. Wang et al., 2014), NTC binding was associated with increases in RBPJ ChIP-Seq signals and H3K27ac signals at promoter and enhancer sites (Figure 3C). Motif analysis revealed that the most common motif lying within 300 bp of NTC ChIP-Seq signals is that of RBPJ (Figure 3D), and that the motif for AP1, a factor not associated with NTC binding sites in other cell types (Chatr-Aryamontri et al., 2017; Drier et al., 2016; Petrovic et al., 2019; Ryan et al., 2017; H. Wang et al., 2014), is also highly enriched in this 600 bp window. Based on the method of Severson et al. (Severson et al., 2017), approximately 13% of NTC binding sites in IC8 cells are predicted to be sequence paired sites (Figure 3E), a specialized type of response element that binds NTC dimers (Arnett et al., 2010). Finally, particularly at early time points, NTC binding sites were spatially associated with genes that are upregulated by Notch, whereas genes that decreased in expression were no more likely to be associated with NTC binding sites than genes that did not change in expression (Figure 3F). Taken together, these studies show that NTCs mainly bind lineage-specific enhancers in IC8 cells and that their loading leads to rapid “activation” of Notch responsive elements and upregulation of adjacent genes.

**Figure 3.**
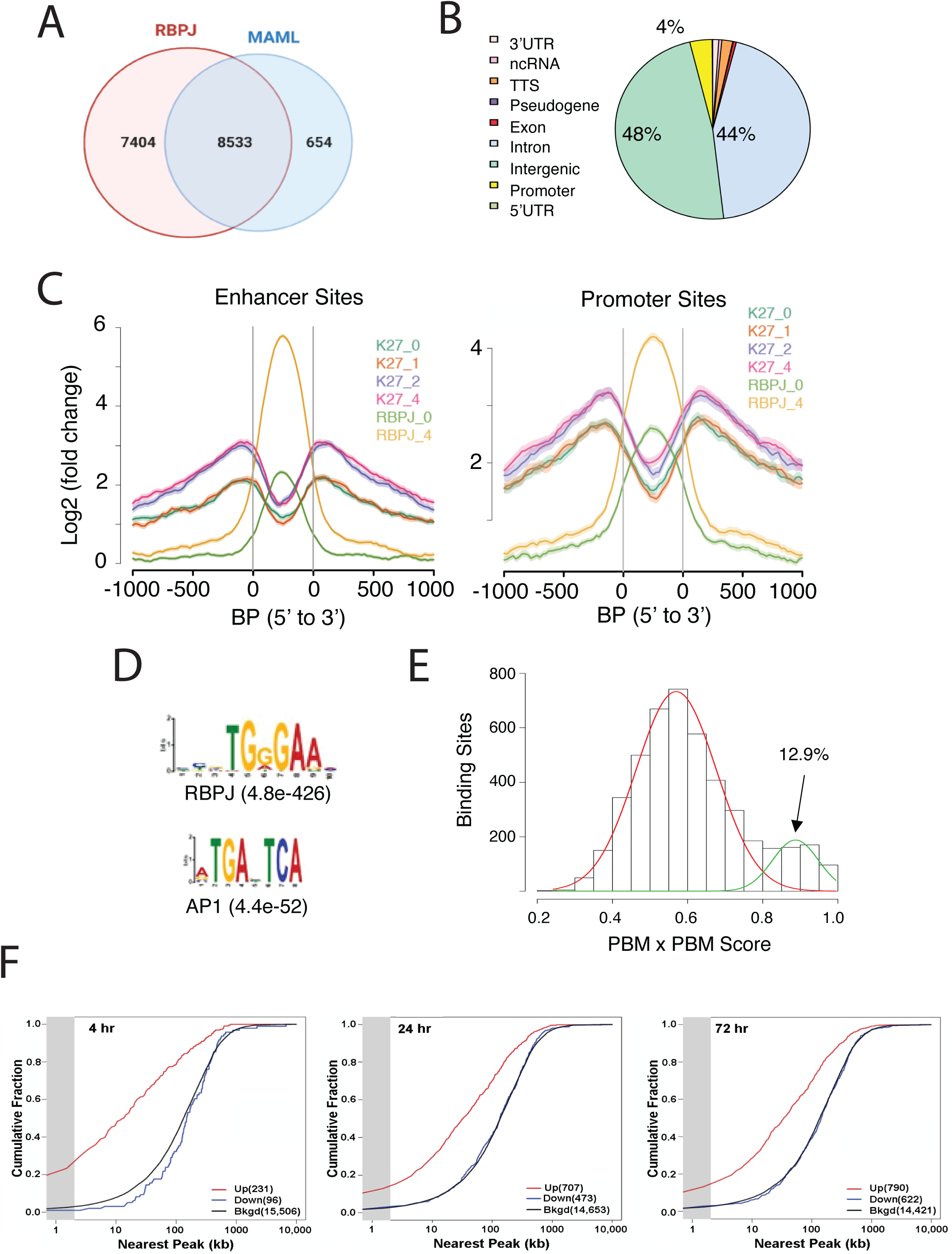
Characterization of Notch transcription complex (NTC) binding sites in IC8-ΔEGF- L1596H cells. (A) Number and overlap of RBPJ and MAML1 binding sites determined by ChIP- Seq of chromatin prepared 4h after Notch activation. (B) Genomic distribution of RBPJ/MAML1 co-binding sites 4h after Notch activation. TTS, transcription termination sites; ncRNA, non- coding RNA. (C) Effect of NTC loading on histone3 lysine27 acetylation (H3K27ac), based on ChIP-Seq for H3K27ac in cells maintained in GSI and in cells 1, 2, and 4h after GSI washout. Transcription factor motifs enriched within 300 bp of RBPJ/MAML1 ChIP-Seq signal peaks. Protein binding matrix (PBM) X PBM scores for NTC binding sites. Sites with scores in the right-hand Gaussian distribution correspond to likely sequence paired sites. (F) Kolmogorov- Smirnov analysis showing spatial relationships between NTC binding sites and transcriptional start sites (TSSs) of genes that increase, decrease, or are unchanged in expression following Notch activation. The gray zone denotes genes with TSSs within 2kb of RBP/MAML1 peaks.

We also performed motif analysis on sites producing significant signals for only RBPJ or only MAML1. RBPJ “only” sites also were enriched for RBPJ (E value 1.3^e-124^) and AP1 (E value 2.8^e-71^) motifs but had lower average ChIP-Seq signals, suggesting these may be weak RBPJ binding sites. MAML1 “only” sites also were enriched for AP1 motifs (E value 2.9^e-181^) but were not associated with RBPJ motifs. These sites were relatively few in number (N=654) and the associated AP1 motifs were distributed broadly around MAML1 signal peaks, arguing against direct physical interaction between MAML1 and AP1 family members on chromatin. Thus, the significance of these “MAML1-only” peaks is uncertain, and it is possible that the observed ChIP-seq signals are non-specific, stemming from over-representation of “open” chromatin in ChIPs.

### *IER5* is a direct Notch target gene

We were intrigued by the convergence of Notch target genes and genes linked to DNA damage/cell stress responses. To test the idea that DNA damage/cell stress response genes contribute to Notch-induced differentiation of squamous cells, we elected to study the gene *IER5* in detail. *IER5* is a member of the immediate early response gene family that encodes a 327 amino acid protein with a ∼50 amino acid N-terminal IER domain and an C-terminal domain predicted to be unstructured. Previous studies have implicated *IER5* in cellular responses to DNA damaging agents and heat shock (Ding et al., 2009; Ishikawa & Sakurai, 2015; Kis et al., 2006), but a role in regulation of keratinocyte growth and differentiation has not been described. Inspection of chromatin landscapes around *IER5* in IC8 cells revealed a series of flanking enhancers, two of which (D and E) showed the largest RBPJ/MAML1 signals and the greatest increase in H3K27ac following Notch activation (Figure 4A). Notably, a similar enhancer landscape exists in non-transformed human keratinocytes (Figure 4, Supplemental Figure 1A), and expression of *IER5* transcripts is readily detectable in normal human skin (Figure 4, Supplemental Figure 1B-D), consistent with the idea that the observed enhancers are involved in physiologic regulation of *IER5* in keratinocytes. Reporter gene assays with enhancers D and E in IC8-ΔEGF-L1596H cells (Figure 4B and 4C, respectively) confirmed that the Notch responsiveness of these elements depend on RBPJ binding sites and also showed, in the case of enhancer D, that a flanking AP1 consensus site is also required. To determine the contributions of enhancers D and E within the genomic *IER5* locus, we used CRISPR/Cas9 targeting to delete the regions containing RBPJ binding sites in these two enhancers in SC2 cells (Figure 4D). These deletions partially abrogated the Notch-dependent increase *IER5* transcription (Figure 4E) and suppressed the accumulation of IER5 protein following Notch activation (Figure 4F), confirming that *IER5* is directly regulated by Notch through these elements.

**Figure 4.**
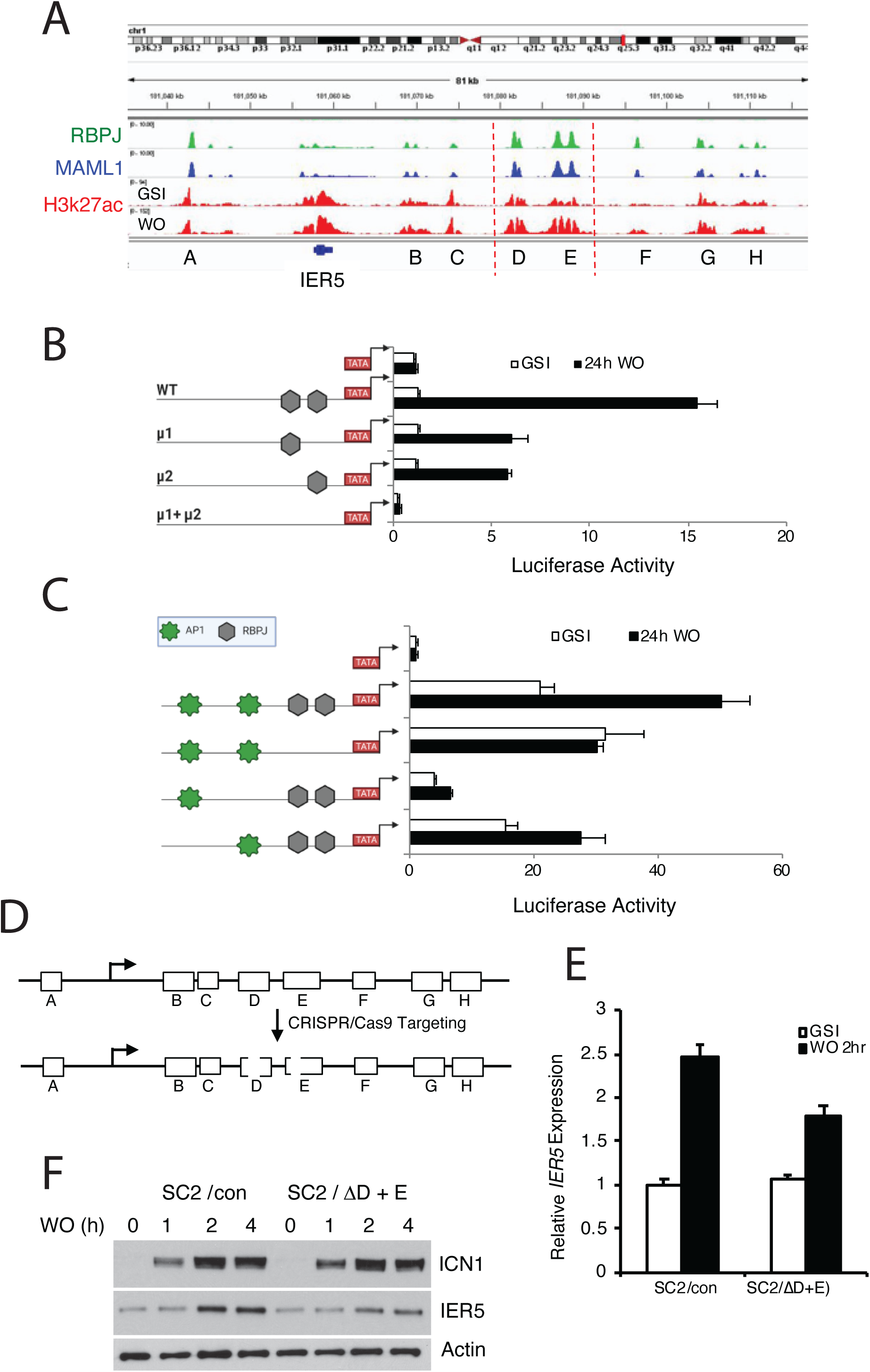
*IER5* is a direct Notch target gene. (A) Chromatin landscapes around *IER5* in IC8- ΔEGF-L1596H cells. ChIP-Seq signals for RBPJ, MAML1, and H3K27ac for cells maintained in and 4h after GSI washout (WO) are shown. (B, C) Activities of a WT *IER5* enhancer E luciferase reporter gene and derivatives bearing mutations (μ) in two RBPJ consensus motifs (B) and a WT *IER5* enhancer D luciferase reporter gene and derivatives bearing mutations in two RBPJ consensus motifs or in flanking AP1 consensus motifs (C). Reporter gene assays were performed in SC2 cells maintained in GSI or 24h after GSI washout (WO). Luciferase reporter gene activity was determined in experimental triplicates and normalized to the activity of a *Renilla* luciferase internal control gene. Error bars represent standard deviations. D) Cartoon showing the CRISPR/Cas9 targeting strategy for *IER5* enhancers D and E. E) Relative *IER5* transcript levels in SC2 cells targeted with control AAVS1 CRISPR/Cas9 plasmids (SC2/con) or with CRISPR/Cas9 plasmids that remove the RPBJ sites in enhancers D and E (SC2/ΔD+E). Cells were either maintained in GSI or were harvested 2h following GSI washout (WO). Transcript abundance was measured in experimental triplicates by RT-PCR and normalized against GAPDH. Error bars represent standard errors of the mean. F) Western blots showing IER5 protein levels in SC2/con cells and SC2/ΔD+E cells that were either maintained in GSI or harvested 1, 2, or 4 h following GSI washout (WO).

### IER5 is required for “late” Notch-dependent differentiation events in squamous cells

To systematically determine the contribution of *IER5* to Notch-dependent changes in gene expression, we compared the transcriptional response to Notch activation in SC2 cells, SC2 cells in which *IER5* was knocked out (I5 cells), and I5 cells to which *IER5* expression was restored (I5AB cells, Figure 5A). Different doses of *IER5* had no effect on gene expression in the absence of Notch signaling, or on the expression of genes that are induced by Notch within 4h; however, by 24h and 72h of Notch activation, I5 cells failed to upregulate a large group of Notch responsive genes that were rescued by add-back of IER5 (Figure 5B, denoted with red box). GO analysis revealed that *IER5*-dependent genes were associated with various aspects of keratinocyte differentiation and biology (Figure 5C; summarized in Tables S6 and S7). An example of a differentiation-associated gene impacted by loss of *IER5* is *KRT1*, a marker of spinous differentiation, expression of which is markedly impaired by *IER5* knockout and restored by *IER5* add-back (Figure 5D). Similarly, *IER5* was required for Notch-dependent expression of the late marker involucrin in 3D cultures (Figure 5G). Thus, *IER5* is necessary but not sufficient for expression of a group of genes that respond to Notch with delayed kinetics.

**Figure 5.**
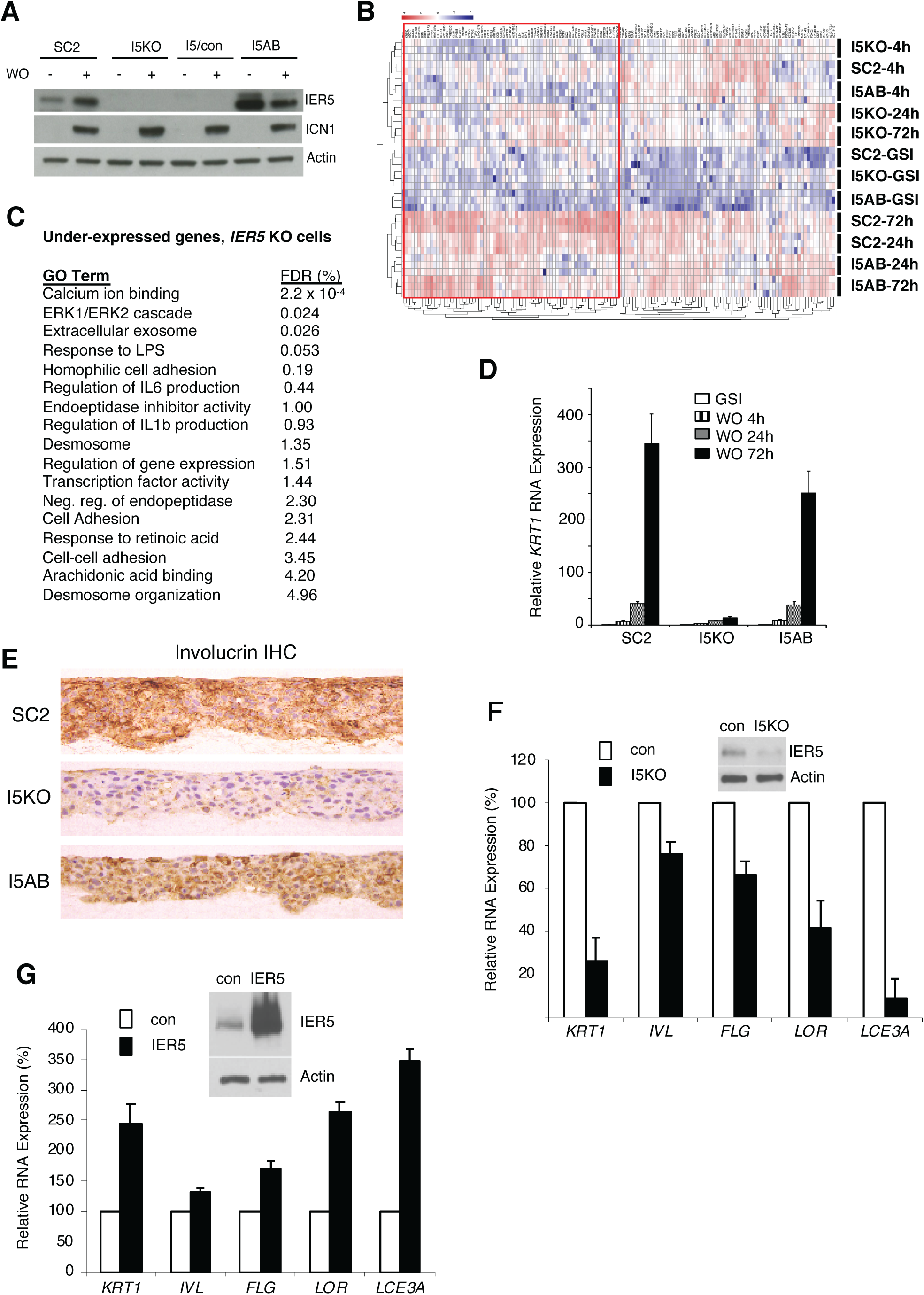
Effect of *IER5* on Notch-dependent changes in gene expression in SC2 cells and NOK1 cells. (A) Western blots showing IER5 and ICN1 protein levels in SC2 cells, a single cell clone derived from SC2-*IER5* knockout cells (I5KO), pooled I5KO cells transduced with empty virus (I5/con), and pooled I5KO cells transduced with *IER5* cDNA (I5AB) that were maintained in GSI (-) or harvested 48h after GSI washout (+). (B) Heat map showing Notch-induced changes in gene expression in SC2 cells, I5KO cells, and I5AB cells. RNA-seq was performed in triplicate at time 0, 4h, 24h, and 72h after GSI washout. Samples were subjected to unsupervised clustering using a gene set containing all genes that were significantly upregulated at any time point after Notch activation in SC2 cells. The red box highlights genes that are under-expressed in *IER5* knockout cells (I5KO) and rescued by re-expression of *IER5* (I5AB). (C) Clustered gene ontogeny (GO) terms associated with the set of under-expressed genes in I5KO cells following Notch activation; see Tables S6 and S7 for a complete list of unclustered and clustered GO terms, respectively. FDR = false discovery rate. (D) Induction of *KRT1* expression following Notch activation in SC2 cells, I5KO cells and I5AB cells. E) Immunohistochemical staining for involucrin in SC2, I5KO, and I5AB cells in raft cultures grown in the Notch-on state (in the absence of GSI). (F, G) Effect of CRISPR/Cas9 targeting of *IER5* and enforced *IER5* expression on differentiation-associated transcripts in NOK1 cells. In F, NOK1 cells were transfected with CRISPR/Cas9 and *IER5* (I5KO) gRNA or AAVS1 control (con) gRNA. In G, NOK1 cells were transduced with empty retrovirus (con) or *IER5* cDNA. In F and G, analyses were done on pooled transfectants and transductants, respectively, which were maintained in low Ca2+ medium or moved to high Ca2+ medium for 3 days (F) or 5 days (G) prior to harvest. Inset Western blots show the extent of *IER5* loss (F) and *IER5* overexpression (G) relative to control cells. In D, F, and G, transcript abundance was measured in experimental triplicates by RT-PCR and normalized against GAPDH. Error bars represent standard deviations of the mean.

To determine if *IER5* has similar effects in non-transformed keratinocytes, we performed studies in a TERT-immortalized oral keratinocyte cell line (NOK1) that differentiates following transfer to high Ca^2+^ medium. We observed that IER5 protein levels increased in differentiation medium in a GSI-sensitive fashion (Figure 5, Supplemental Figure 1A) and that transcript levels of *IER5* and the canonical Notch target gene *NRARP* were depressed by GSI (Figure 5, Supplemental Figure 1B, C), consistent with Notch-dependent regulation of *IER5*. Unexpectedly, we did not observed activation of NOTCH1 in NOK1 cells (not shown); instead, differentiation was accompanied by activation of NOTCH2 and NOTCH3 (inferred from the accumulation of smaller polypeptides consistent with ADAM cleavage products under differentiation conditions in the presence of GSI; Figure 5, Supplemental Figure 1A), as well as increased expression of NOTCH3, a known target of activated Notch. Suppression of *IER5* transcript levels by GSI was partial, suggesting that additional pathways influence *IER5* expression, consistent with the complex enhancer landscape around this gene. To study the effect of *IER5* on NOK1 cell differentiation, we studied the effects of targeting *IER5* with CRISPR/Cas9 and enforced expression of *IER5* through retroviral transduction. *IER5* knockout suppressed expression of multiple differentiation-associated genes (Figure 5F), whereas overexpression of *IER5* increased expression of each of these markers (Figure 5G). Thus, *IER5* is required for Notch-dependent differentiation of malignant and non-transformed keratinocytes.

### IER5 binds B55*α*/PP2A complexes

The structure of IER5 suggests that it functions through protein-protein interactions. To identify interacting proteins in an unbiased way, we expressed a tagged form of IER5 in *IER5* null I5 cells and performed tandem affinity purification followed by mass spectrometry (Adelmant, Garg, Tavares, Card, & Marto, 2019), which identified the B55*α* regulatory subunit of PP2A and PP2A scaffolding and catalytic subunits as potential interactors (Figure 6A; summarized in Table S8). We confirmed these associations by expressing tagged IER5 in I5 cells and tagged B55*α* in SC2 cells (Figure 6B). Full-length IER5 and the N-terminal IER domain of IER5 co-precipitated endogenous B55*α* in Notch-independent fashion, whereas the C-terminal portion of IER5 did not (Figure 6C). Similarly, tagged B55*α* co-precipitated endogenous IER5 in a fashion that was augmented by Notch activation (Figure 6D), consistent with increased recovery of IER5 due to induction of *IER5* expression by Notch. To confirm that IER5 binds B55*α* directly, we studied the interaction of purified recombinant proteins. IER5 exhibited saturable binding to B55*α*- coated beads (Figure 6E), and additional microscale thermophoresis studies showed that IER5 binds B55*α* with a Kd of approximately 100nM (Figure 6F).

**Figure 6.**
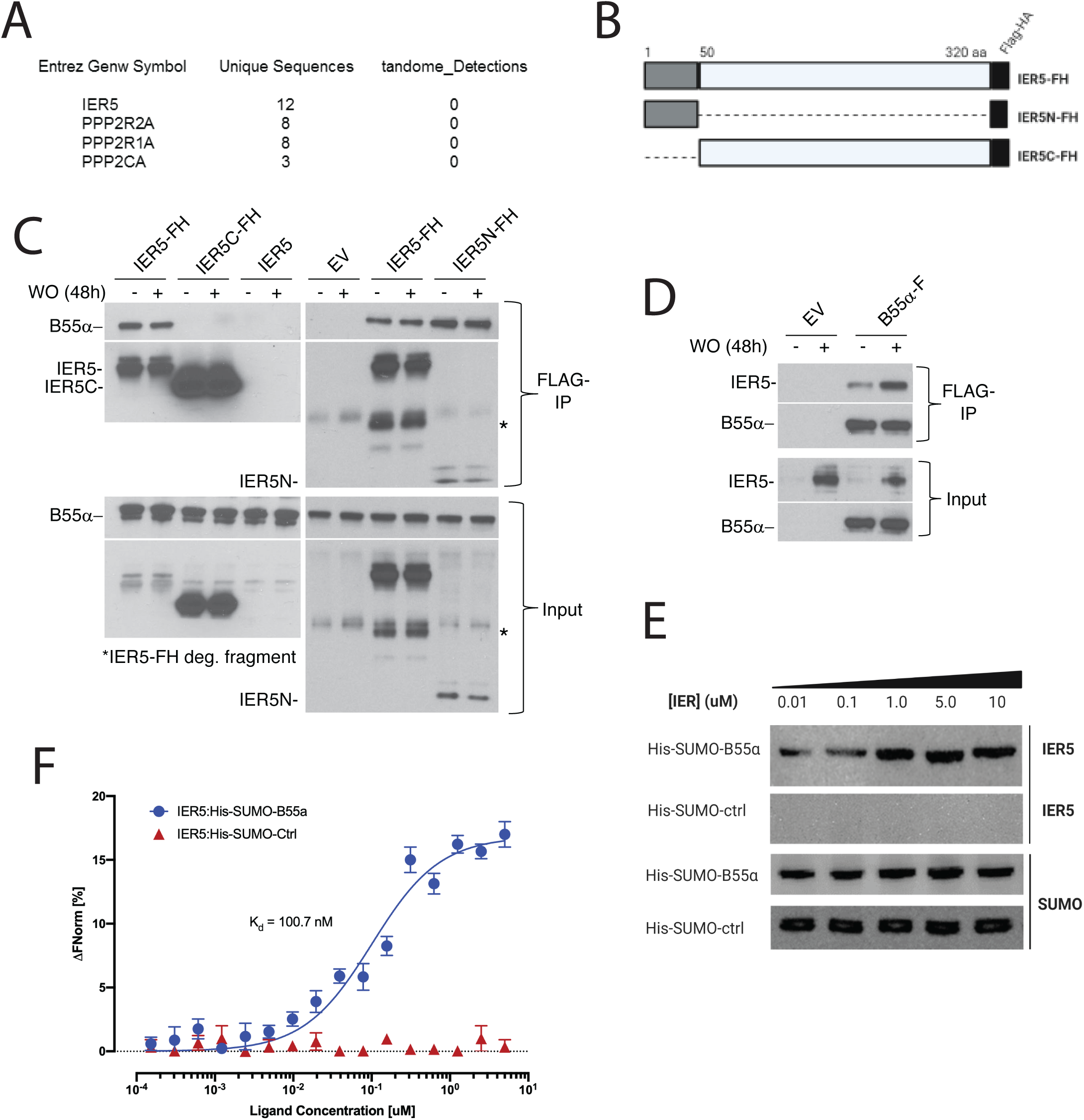
IER5 binds to B55*α*. (A) Polypeptides identified by mass spectroscopy in immunoprecipitates prepared from I5 cells expressing tandem-tagged IER5. (B) Cartoon showing the structure of tandem-tagged IER5 polypeptides. FH, FLAG-HA tag. (C) Western blot analysis of immunoprecipitates prepared from I5 cells expressing the indicated forms of tagged IER5. WO, washout. (D) Western blot analysis of immunoprecipitates prepared from SC2 cells transduced with empty virus (EV) or virus expressing FLAG-tagged B55*α*. (E) Western blot showing that IER5 binds His-Sumo-tagged B55*α* immobilized on beads. The upper two panels were stained for IER5, while the lower two panels were stained for SUMO. (F) Microscale thermophoresis showing saturable binding of IER5 to His-Sumo-tagged B55*α*.

### *IER5* is epistatic to *PPP2R2A* in SCC cells

To gain insight into the role of B55*α* in regulation of Notch- and *IER5*-sensitive genes, we isolated SC2 cell clones that were knocked out for *IER5, PPP2R2A*, or both genes (Figure 7A). Knockout of *PPP2R2A* did not affect the levels of ICN1 following GSI washout (Figure 7A), but increased the expression of *KRT1* in 2D cultures (Figure 7B) and the accumulation of involucrin in 3D cultures (Figure 7C), effects that were suppressed by add-back of B55*α*, suggesting a model in which *IER5* suppresses a B55*α*-dependent activity. This was supported by assays performed with *IER5*/*PPP2R2A* double knockout cells (Figure 7A), in which expression of the late genes such as *KRT1* was restored in the absence of *IER5* (Figure 7D). These results suggest that Notch-mediated suppression of B55*α*-PP2A activity via IER5 is important in regulating the complex series of events downstream of Notch that lead to squamous cell differentiation.

**Figure 7.**
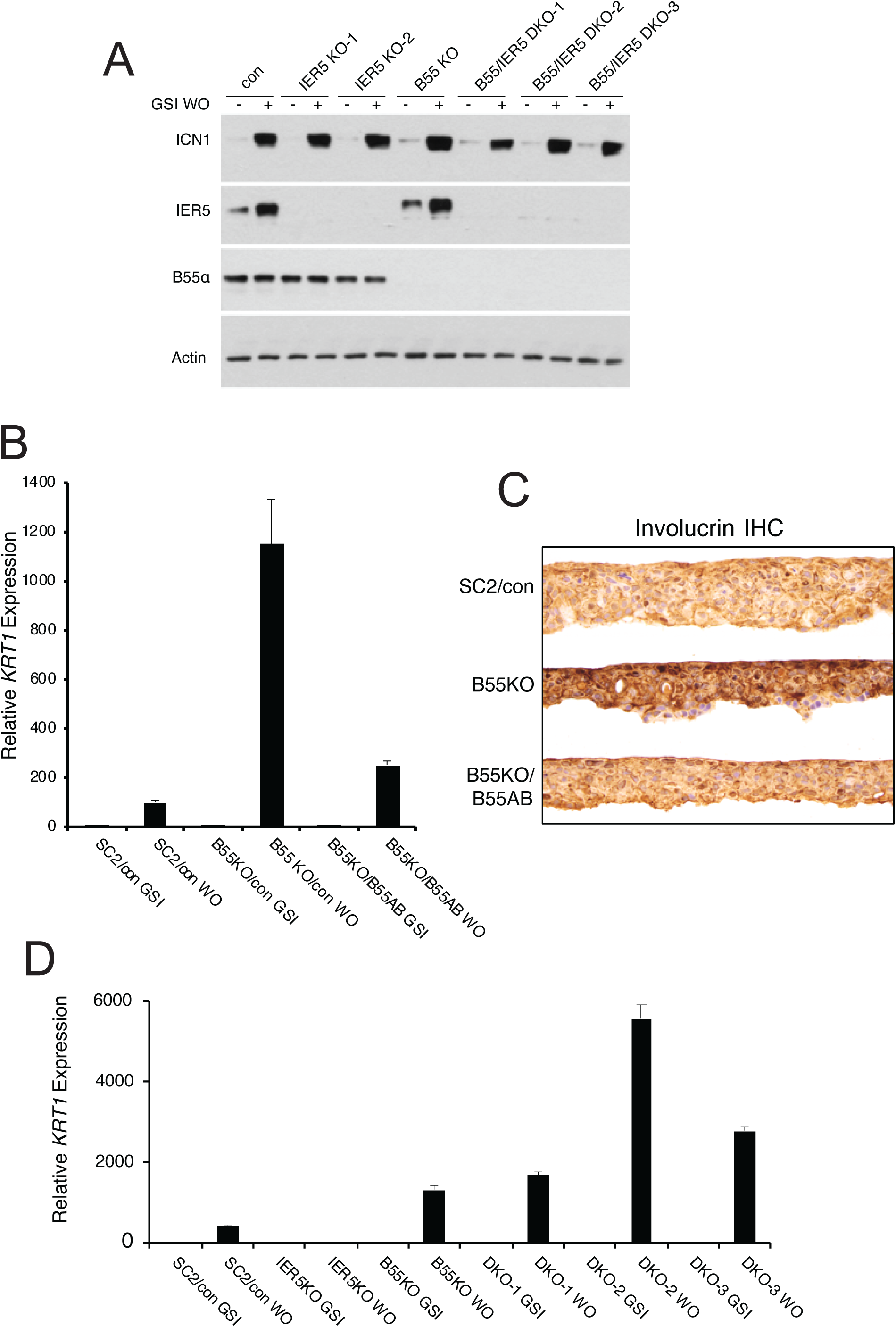
*PPP2R2A* is epistatic to *IER5*. (A) Western blot showing IER5 and B55*α* protein levels in single (KO) and double (DKO) *PPP2R2A* and *IER5* knockout clones in the presence of GSI (-) and 72h after GSI washout (+). (B) *PPP2R2A* knockout enhances Notch-dependent expression of *KRT1*. RT-PCR analysis of *KRT1* expression in SC2 cells transduced with an empty retrovirus (SC2/con); *PPP2R2A* knockout cells (B55 KO) transduced with empty retrovirus (B55KO/con); and *PPP2R2A* knockout cells transduced with B55α-expressing retrovirus (B55KO/B55AB). WO = GSI washout. (C) Immunohistochemical (IHC) staining of SC2 control, B55KO, and B55KO/B55AB cells for involucrin in raft cultures in GSI-free medium. (D) B55*α* knockout negates the requirement for *IER5* for Notch-dependent upregulation of *KRT1*. Results are shown for SC2 control cells (SC2/con); an *IER5* knockout clone; a *PPP2R2A* knockout clone (B55KO); and three *IER5*/*PPP2R2A* double knockout (DKO) clones. Cells were maintained in GSI or harvested 72h following GSI washout (WO). *KRT1* transcript abundance was measured in experimental triplicates by RT-PCR and normalized against GAPDH. Error bars represent standard errors of the mean.

## Discussion

Our work provides a genome-wide view of the direct effects of Notch in squamous cells, in which Notch activation induces growth arrest and differentiation. The phenotypic changes induced by Notch are mediated by a largely squamous cell-specific transcriptional program that includes genes linked to keratinocyte differentiation and DNA damage responses, including *IER5*, which modulates the activity of B55*α*-containing PP2A complexes. Upregulation of these Notch-responsive genes are associated with binding of NTCs to RBPJ sites within lineage-specific enhancers, including a minority of sequence-paired sites, a specialized dimeric NTC binding element recently implicated in anti-parasite immune responses in mice (Kobia et al., 2020). The Notch target genes and elements identified here in malignant squamous cells are likely to be relevant for understanding Notch function in non-transformed squamous cells, as many of the NTC binding enhancers found near Notch target genes in SCC cells are also active in normal human keratinocytes (based on review of ENCODE data for non-transformed human keratinocytes). These observations have a number of implications for understanding how Notch regulates the growth and differentiation of squamous cells and highlight the potential for Notch to influence the activity of diverse signaling pathways in keratinocytes through modulation of PP2A.

Prior work has suggested that p53-mediated upregulation of Notch expression and activity is a component of the DNA damage response in keratinocytes (Mandinova et al., 2008). Conversely, our work shows that Notch activation, even in a *TP53* mutant background, induces the expression of genes that are components of the keratinocyte DNA damage/cell stress response. In cells of most lineages, p53 responses serve to suppress cell growth and, if DNA damage is severe or cannot be repaired, to induce apoptosis. However, our work suggests that in cells of squamous lineage these two pathways share a core set of genes that induce differentiation. In the case of p53, this function may act to help to eliminate damaged cells from the epidermal stem cell pool without inducing apoptosis, which could comprise the barrier function of squamous epithelial surfaces. Consistent with this idea, recent work has shown that low-dose radiation induces the differentiation of esophageal squamous cells at the expense of self-renewal *in vivo* (Fernandez-Antoran et al., 2019). *TP53* and Notch genes are frequently co-mutated in squamous carcinomas of the skin (as in our model system) and other sites; although co-mutation of tumor suppressor genes in a particular cancer is typically taken as evidence of complementary anti-oncogenic activities, our work suggests that in the cases of squamous carcinoma it may reflect, at least in part, redundant Notch and p53 functions. Further work delineating the crosstalk between p53 and Notch signaling in well controlled model systems will be needed to further test this idea.

Among the genes linked to Notch and p53 is *IER5*, an immediate early response gene that is a component of the DNA-damage response in a number of cell types (Ding et al., 2009; Kis et al., 2006; Kumar et al., 1998; Yang et al., 2016). Several lines of investigation suggest that IER5 modulates the function of PP2A complexes containing B55 (Asano et al., 2016; Ishikawa, Kawabata, & Sakurai, 2015; Kawabata, Ishita, Ishikawa, & Sakurai, 2015) regulatory subunits, and here we demonstrate that IER5 directly binds B55*α* protein in a purified system. However, the exact effect of IER5 on PP2A function is uncertain. One model suggests that IER5 augments the ability of B55/PP2A complexes to recognize and dephosphorylate specific substrates such as S6 kinase and HSF1 (Ishikawa et al., 2015; Kawabata et al., 2015), the latter leading to HSF1 activation as part of the heat shock response. However, our work with double knockout cells suggests IER5 inhibits at least some activities that are attributable to B55*α*-containing PP2A complexes. Further work in purified systems may be helpful in clarifying how IER5 influences B55/PP2A function. We also note that while IER5 is necessary for upregulation of genes that are induced by Notch with delayed kinetics, it is not sufficient; therefore, it is likely that other PP2A- regulated factors that work in concert with Notch to induce expression of this class of genes await discovery.

Finally, we note that our small screen of SCC cell lines suggests that squamous cell carcinomas retain the capacity to respond to Notch signals by undergoing growth arrest and differentiation. Although originally identified as an oncogene, sequencing of cancer genomes has revealed that Notch most commonly acts as a tumor suppressor, particularly in squamous cell carcinoma, which is difficult to treat when advanced in stage. While restoring the expression of defective Notch receptors is challenging, detailed analysis of crosstalk between Notch and other pathways may reveal druggable targets leading to reactivation of tumor suppressive signaling nodes downstream of Notch, which would constitute a new therapeutic strategy for squamous cancers.

## Materials and Methods

### Cell lines and 2D cultures

Cells were grown under 5% C0_2_ at 37° C in media supplemented with glutamine and streptomycin/penicillin. IC8 cells (N. J. Wang et al., 2011) were cultured in Keratinocyte medium as described (Purdie, Pourreyron, & South, 2011). 7EGF-L1596H cDNA cloned into pBABE-puro was packaged into pseudotyped retrovirus and used to transduce IC8 and SCCT2 cells, which were selected with puromycin (1 ug/ml). In some instances, cells were also transduced with pseudotyped MigRI retrovirus encoding dominant negative MAML1 fused to GFP (Weng et al., 2003). IC8 cell clones were isolated by single cell sorting in the Dana Farber Cancer Institute Flow Cytometry Core Facility. NOK1 cells were grown in keratinocyte-SFM medium supplemented with human EGF and bovine pituitary extract (BPE) (Thermo Fisher Scientific) and induced to differentiate by transfer to Dulbcecco modified Eagle medium (DMEM) containing 10% fetal bovine serum. Cells were carried in the presence of the GSI compound E (*N*-[(1*S*)-2-[[(3*S*)-2,3-Dihydro-1-methyl-2-oxo-5-phenyl-1*H*-1,4-benzodiazepin-3- yl]amino]-1-methyl-2-oxoethyl]-3,5-difluorobenzeneacetamide; Tocris), which was used at 1μM in all experiments. Washout controls cells were treated with vehicle (0.1% dimethylsufoxide, DMSO).

### Cell growth assays

Cell numbers were estimated using CellTiter Blue (Promega) per the manufacturer’s recommendations. Fluorescence was measured using a SpectraMax M3 microplate reader (Molecular Devices).

### Organotypic 3D cultures

3D raft cultures were performed on a matrix containing 5×10^5^ J2 3T3 fibroblast cells and rat collagen as described (Arnette, Koetsier, Hoover, Getsios, & Green, 2016). Briefly, rafts were allowed to mature for 6-7 days and then were seeded with 5×10^5^ SCC cells in E-medium with or without GSI. After 2 days, rafts were raised to the fluid-air interface, and medium was refreshed every 2 days for a total of 12 additional days.

### Targeted exon sequencing

NGS was performed on IC8 and SCCT2 cell genomic DNA using the “oncopanel” assay (Abo et al., 2015; Wagle et al., 2012), which covers 447 cancer genes. Briefly, DNA (200 ng) was enriched with the Agilent SureSelect hybrid capture kit and used for library preparation. Following sequencing (Illumina HiSeq 2500), reads were aligned to human genome GRCh37 (hg19) (Li & Durbin, 2009), sorted, duplicate marked, and indexed. Base-quality score calibration and alignments around indels was done with Genome Analysis Toolkit (DePristo et al., 2011; McKenna et al., 2010). Single nucleotide variant calls were with MuTect (Cibulskis et al., 2013). Copy number alterations were determined using RobustCNV. Structural variants were detected using BreaKmer (Abo et al., 2015).

### Preparation of ChIP-Seq and RNA-seq Libraries

Chromatin was prepared as described (H. Wang et al., 2014) and was immunoprecipitated with antibodies against MAML1 (clone D3K7B) or RBPJ (clone D10A11, both from Cell Signaling Technology) and Dynabeads bearing sheep anti-rabbit Ig (Thermo Fisher Scientific). H3K27ac ChIPs were prepared using the ChIP assay kit (Millipore) and H2K27ac antibody (ab4729, Abcam). ChIP-seq libraries were constructed using the NEBNext Ultra II DNA Library Prep Kit (New England BioLabs). Total RNA was prepared with Trizol (Life Technology) and RNeasy Mini columns (Qiagen). RNA libraries were constructed using the NEBNext Ultra II RNA Library Prep kit (New England BioLab). ChIP-seq and RNA-seq libraries were sequenced on an Illumina NextSeq 500 instrument. ChIP-seq and RNA-seq data sets will be deposited in GEO.

### ChIP-seq data analysis

Reads were trimmed with Trim Galore (v.0.3.7 using cutadapt v.1.8), assessed for quality with FastQC (v.0.11.3), and aligned to GRCh38/hg38 with bowtie (v.2.0.0) (Langmead & Salzberg, 2012). Peaks were identified using MACS2 (v.2.1.1 (Zhang et al., 2008)) and annotated using Homer (v3.12, 6-8-2012 (Heinz et al., 2010)). Peaks mapping to repeats (repeatMasker track, from UCSC) or ENCODE blacklisted regions were removed. Overlapping RBPJ and MAML1 peaks were identified with bedtools intersectBed (v2.23.0) (Quinlan & Hall, 2010). Motif analysis was performed using MEME-ChIP (Machanick & Bailey, 2011). Average signal profiles were generated with ngsplot (Shen, Shao, Liu, & Nestler, 2014). RBPJ sequence-paired sites (SPSs) were identified as described (Severson et al., 2017). Mixed Gaussian curves were generated in R using the mixtools (v.1.1.0) function (Benaglia, Chauveau, Hunter, & Young, 2009).

### RNA-seq data analysis

Reads were trimmed as described for DNA reads and aligned to human genome GRCh38/hg38 using gencode release 27 annotations and STAR (v.2.5.3a) (Dobin et al., 2013). Raw counts from two sequencing runs were loaded into R (Team, 2014), summed, and filtered to exclude transcripts with <0.5 reads per million mapped before performing differential expression (DE) analysis with edgeR (v.3.16.5) and RUVSeq (v.1.8.0) (Risso, Ngai, Speed, & Dudoit, 2014). After first-pass DE analysis, a control set of 733 genes with FDR>0.5 in all pairwise comparisons was used with RUVg (k=1) to identify unwanted variation. Second-pass edgeR analysis included the RUVg weights in the model matrix. Genes with FDR<5% and absolute logFC >1 were retained for further analysis. DAVID v.6.8 (Huang da, Sherman, & Lempicki, 2009) was used for gene ontology (GO) enrichment analysis of the DE gene lists. EdgeR cpm function with library size normalization and log2 conversion was used to generate expression values, which were displayed using pheatmap (R package version 1.0.8). Other plots were made using in-house R scripts (available upon request).

### Quantitative RT-PCR

Total RNA was isolated using RNAeasy Mini Kit (Qiagen) and cDNA was prepared using an iScript cDNA synthesis kit (BioRad). qPCR was carried out using a CFX384 Real-Time PCR Detection System. Gene expression was normalized to GAPDH using the *Δ*Δ CT method. Primer sets used are available on request.

### In Situ Hybridization

*In situ* hybridization (ISH) reagents were from Advanced Cell Diagnostics. Deidentified normal human skin was obtained from the paraffin archives of the Department of Pathology at Brigham and Women’s Hospital under institutional review board protocol #2014P001256. Briefly, 4μ sections of skin were deparaffinized, processed using RNAscope 2.5 Universal Pretreatment Reagent, and hybridized to probes specific for human *IER5*, human *PPIB* (peptidylprolyl isomerase B), or bacterial DapB in a HybEZ II oven. ISH signal was developed using the RNAscope 2.5 HD Assay.

### Immunostaining of cells and organotypic rafts

For indirect immunofluorescence microscopy, cells grown on chamber microscope slides were fixed in 4% paraformaldehyde and treated with immunofluorescence blocking buffer (catalog #12411, Cell Signaling Technology). Staining with primary antibodies against involucrin (Sigma-Aldrich, catalog #I9018, 1:500) or plakophilin-1 (Sigma-Aldrich, catalog #HPA027221, 1:300) in antibody dilution buffer (catalog #12378, Cell Signaling Technology) was developed by incubation with Alexa Fluor-conjugated secondary antibodies (Cell Signaling Technology, 1:1000). After counterstaining with DAPI (BioLegend, catalog #422801), slides were coverslipped with ProLong Gold Antifade Reagent (Cell Signaling Technology, catalog #9071) and imaged on a Nikon 80i immunofluorescence microscope. Rafts were fixed in 4% buffered formalin for 24h, processed, and embedded in paraffin. Sections (4μ) were placed on Superfrost Plus slides and baked at 60°C for 1h. Immunohistochemical staining was performed on a Leica Bond III instrument using the following primary antibodies and Leica antigen retrieval conditions: involucrin (Sigma-Aldrich, catalog #I9018, 1:10,000), retrieval H1 (30min); plakophilin1 (Sigma-Aldrich, catalog #HPA027221, 1:500), retrieval H1 (30min); keratin-1 (Abcam, catalog #185628, 1:1,000), H1 retrieval (30min); Ki67 (BioCare, catalog# CRM325, 1:100), retrieval H2 (20min); or ICN1 (Cell Signaling Technology, catalog #4147, 1:50), retrieval H2 (40min). Diaminobenzidine (DAB) staining was developed using the Bond Polymer Refine Detection Kit (Leica). Slides were counterstained with hematoxylin. Digital micrographs were captured with an Olympus BX40 microscope and Olympus cellSens Entry software.

### Reporter gene assays

Luciferase reporter genes containing *IER5*-associated enhancers were assembled in pGL3-TATA (H. Wang et al., 2014). Mutatagenesis was with the QuickChange II kit (Agilent Technologies). Luciferase assays were performed using Dual Luciferase Assay Kit (Promega) as described (Malecki et al., 2006) using lysates from cultured cells that were co-transfected with firefly luciferase and internal control *Renilla* luciferase plasmids using Lipofactamine 2000 (Thermo Fisher Scientific).

### Western blotting and immunoprecipitation

Whole cell lysates were prepared as described (Malecki et al., 2006). Protein concentration was measured by Bradford assay (Bio-Rad) prior to SDS-PAGE. Western blots were stained with the following antibodies: ICN1 (clone D3B8), B55α (clone 2G9), NOTCH2 (clone D76A6), NOTCH3 (D11B8), ID3 (clone D16D10), and GADD45A (clone D17E8) (all from Cell Signaling Technology); SUMO (Lifesensors, catalog# AB7002); and IER5 (catalog# HPA029894), involucrin (catalog# I9018), FLAG epitope (catalog# F3165), and actin (catalog# A1978) (all from Sigma). Secondary antibodies were goat anti-rabbit (catalog# 7074) or horse anti-mouse (catalog# 7076) IgG (Cell Signaling Technology), or goat anti-chicken IgY (Abcam), all conjugated to horseradish peroxidase. Staining was developed with Super Signal West Pico Chemiluminescent Substrate (Thermo Scientific). To prepare immunoprecipitates, cells were lysed in 50 mM Tris, pH 7.4, containing 150 mM NaCl, 1mM EDTA, 1% Triton X-100, and protease inhibitors (Sigma). Lysates were incubated overnight at 4°C with 20μl anti-FLAG M2 magnetic beads (Sigma). After extensive washing, bound proteins were eluted with FLAG peptide (Sigma) and analyzed on Western blots as above.

### Tandem affinity purification of IER5 complexes and mass spectrometry

Mass spectroscopy was performed on tandem affinity purified IER5 complexes prepared from IE5 knockout (I5) cells expressing tandem tagged IER5 48h after GSI washout. Lysis of cells and subsequent tandem purification were as described (Adelmant et al., 2019). Tryptic peptides were analyzed by electrospray mass spectrometry (QExactive HF mass spectrometer, Thermo Fisher; Digital PicoView electrospray source platform, New Objective). Spectra were recalibrated using the background ion (Si(CH3)2O)6 at m/z 445.12 +/- 0.03 and converted to a Mascot generic file format (.mgf) using multiplierz scripts (Askenazi, Parikh, & Marto, 2009; Parikh et al., 2009). Spectra were searched using Mascot (v2.6) against three databases: i) human protein sequences (downloaded from RefSeq); ii) common lab contaminants; and iii) a decoy database generated by reversing the sequences from these two databases. Spectra matching peptides from the reverse database were used to calculate a global FDR and were discarded, as were matches to the forward database with FDR>1.0% and those present in >1% of 108 negative tandem affinity purification controls (Rozenblatt-Rosen et al., 2012).

### Recombinant protein expression and purification

A cDNA encoding B55*α* with 6xHis-SUMO N-terminal tag was cloned into the baculovirus transfer vector pVL1392. High-titer baculovirus supernatants were used to infect insect Sf9 cells grown at a density of 4.0×10^6^ cell/mL. After 72h of incubation at 27°C, conditioned media was isolated by centrifugation, supplemented with 20mM Tris buffer, pH 7.5, containing 150mM NaCl, 5mM CaCl_2_, 1mM NiCl_2_, and 0.01mM ZnCl_2_, re-centrifuged to remove residual debris, and applied to a Ni-NTA column. After washing with 20mM Tris buffer, pH 7.5, containing 150mM NaCl and 5 mMCaCl_2_, B55*α* was eluted in the same buffer supplemented with 500mM imidazole. This eluate was concentrated with a centrifugal filter and then subject to S200 size exclusion chromatography in 20mM Na cacodylate, pH 6.0, containing 150mM NaCl, and 5mM CaCl_2_. Fractions containing B55*α* were further purified by ion exchange chromatography on a MonoQ column in 20 mM Na cacodylate, pH 6.0, using a linear NaCl gradient. The purified protein was buffer-exchanged into 20mM HEPES buffer, pH 7.5, containing 150mM NaCl, prior to flash freezing and storage at −80°C. A cDNA encoding IER5 with a N-terminal 6xHis-SUMO tag was cloned into pTD6 and used to transform Rosetta *E. coli* cells. Expression of IER5 was induced at 37°C for 4h with IPTG induction followed by inclusion body preparation. Lysates were centrifuged at 4°C for 20min at 30,000 x g and pellets were resuspended in 10mL wash buffer (50mM Tris-HCl, pH 7.5, 150mM NaCl, containing 1% Triton X-100 and 1M urea) per gram cell weight, and incubated at 23°C for 5min. Following multiple washes, inclusion bodies were resuspended in extraction buffer (50mM Tris-HCl, pH 7.5, 8M urea, 1mM *β*- mercaptoethanol, 1mM PMSF) and incubated at room temperature for 1h. The solubilized proteins were then dialyzed overnight against a 100-fold volume of wash buffer and cleared by centrifugation. 6x-his-IER5 was then concentrated on Ni-NTA beads, eluted with 500mM imidazole cleaved using SUMO protease (SUMOpro, Lifesensors), and passed back over Ni-NTA beads. Untagged IER5 was then purified by chromatography on S200 and MonoQ columns as described for B55*α*.

### IER5/B55*α* binding assays

For bead pulldown assays, 5μM Purified His-SUMO-B55*α* bait was bound to 100ul Ni-NTA beads, which were was and mixed with different concentrations of purified IER5 in 20mM HEPES buffer, pH 7.5, containing 150mM NaCl for 1h. Control binding assays were conducted with purified His-SUMO bond to Ni-NTA beads. Following extensive washing, proteins were eluted by boiling in SDS-PAGE loading buffer and analyzed on SDS-PAGE gels followed by Western blotting. Microscale thermophoresis assays were performed on a NanoTemper ® Monolith NT.115 instrument with blue/red filters (NanoTemper Technologies GmbH, Munich, Germany). Samples were prepared in 20mM Tris-HCl, pH 7.5, containing 150mM NaCl, 1mM *β*-mercaptoethanol, 1% glycerol, 0.05% Tween-20, and loaded into premium treated capillaries. Measurements were performed at 22° C using 20% MST power with laser off/on times of 5s and 30s, respectively. His-SUMO-B55a target labeled with red fluorescent detector as per the NanoTemper His-labeling kit was used at a concentration of 10nM and mixed with 16 serial dilutions of purified IER5 ligand from 5μM to 0.000153μM. All experiments were repeated three times. Data analyses were performed using NanoTemper® analysis software.

### CRISPR/Cas9 targeting and site directed mutagenesis

Guide RNAs (gRNAs) were designed using software available at http://crispr.mit.edu. To score the effects of gene editing in bulk cell populations, gRNAs were cloned into pL-CRISPR.SFFV.GFP (Addgene) and transiently transfected using Lipofactamine 2000 (Thermo Fisher Scientific). Cells sorted for GFP expression 48h post-transfection were used in downstream analyses. To create double knockout cells, single cell clones bearing single gene knockouts were transduced with pL-CRISPR.SFFV.GFP bearing gRNA. Lentivirus was packaged by co-transfection with psPAX2 and pMD2.G. To create deletions, pairs of gRNAs flanking genomic regions of interest were cloned into lentiCRISPRv2 neo (addgene) and lentiCRISPRv2 hygro (addgene), or into pL-CRISPR.SFFV.GFP and pL-CRISPR.SFFV.RFP (Addgene). Double deletants were isolated sequentially by selection for RFP/GFP double positivity and G418/hygromycin resistance. The sequences of gRNAs used are available on request.

## Supporting information

Differentially expressed genes after 4h of Notch activation, SC2 cells.

Differentially expressed genes after 24h of Notch activation, SC2 cells.

Differentially expressed genes after 72h of Notch activation, SC2 cells.

Unclustered GO annotations of Notch-sensitive genes, SC2 cells, at 72h of Notch activation.

Clustered GO annotations of Notch-sensitive genes, SC2 cells, at 72h of Notch activation.

Unclustered GO annotations of IER5-dependent Notch-sensitive genes, SC2 cells.

Clustered GO annotations of IER5-dependent Notch-sensitive genes, SC2 cells.

IER5-interacting proteins identified by tandem affinity purification.

## Figure Legends

**Figure 1, Supplemental Figure 1.**
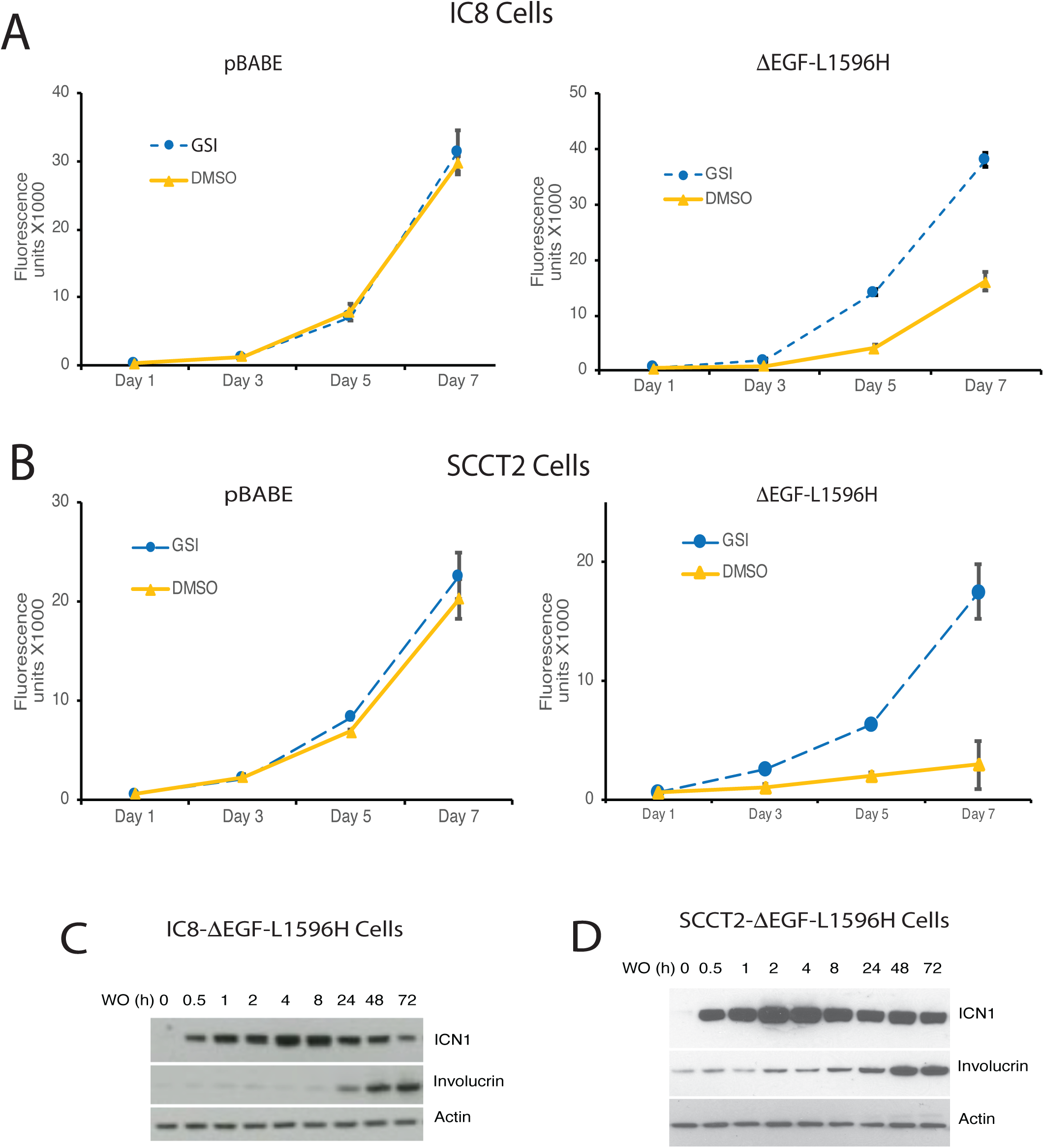
Notch activation induces differentiation and growth arrest of the squamous carcinoma cell lines IC8 and SCCT2. (A, B) Effects of GSI and sham washout on the growth of IC8 and SCCT2 cells transduced with either empty vector (pBABE) or ΔEGF- L1596H. Cell numbers at various times post-GSI washout (DMSO) or mock GSI-washout (GSI) were assessed using Cell Titer-Blue on independent cultures in quadruplicate. (C, D) Western blot showing the kinetics of ICN1 generation and increases in involucrin (IVL) following GSI washout (WO) in IC8 and SCCT2 cells transduced with ΔEGF-L1596H.

**Figure 1, Supplemental Figure 2.**
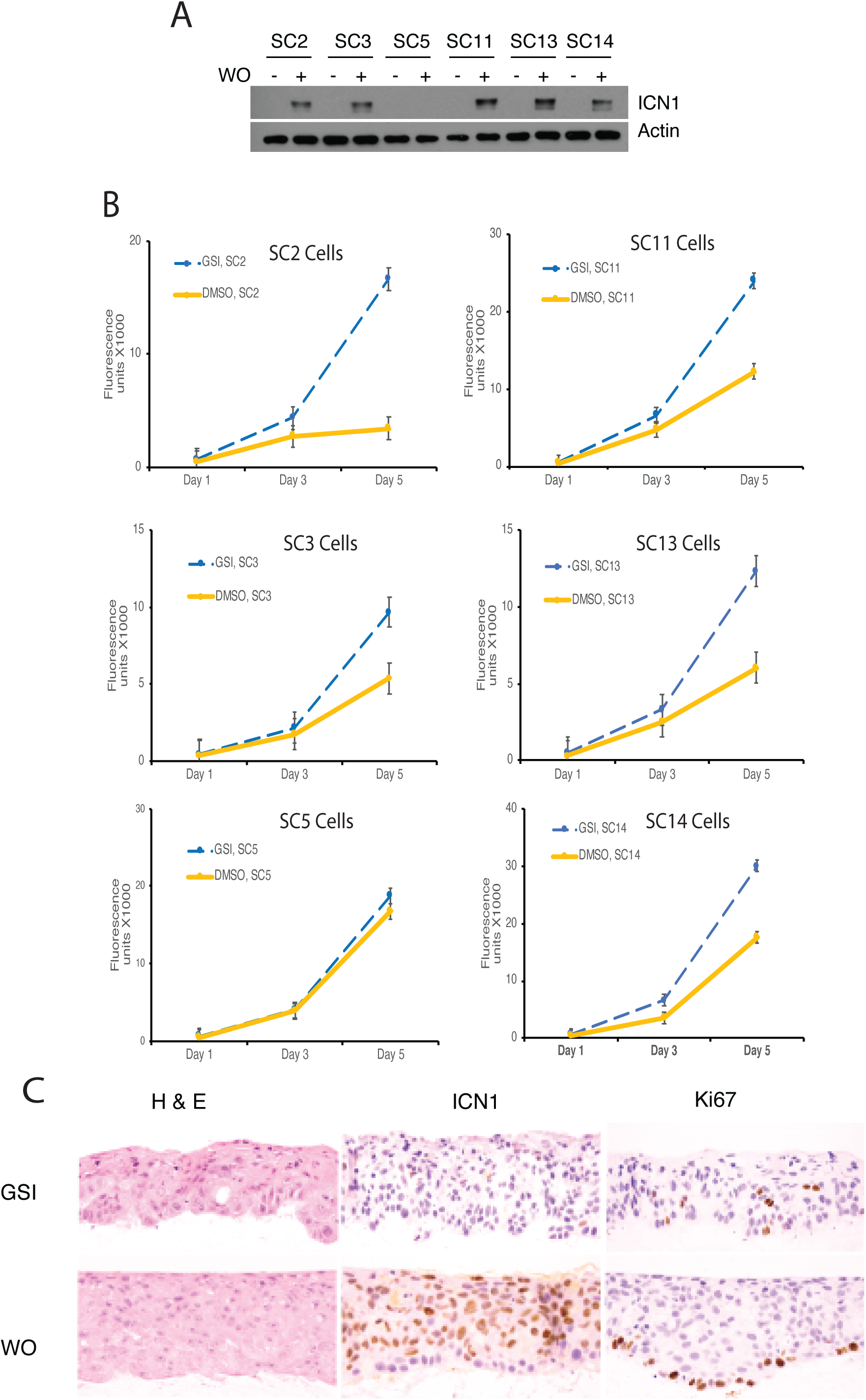
Characterization of clones derived from single IC8 cells transduced with ΔEGF-L1596H. (A) Western blot showing ICN1 levels 4h post-GSI washout in IC8-ΔEGF-L1596H clones. (B) Effects of Notch activation on growth of IC8-ΔEGF-L1596H clones. Cell numbers at various times post-GSI washout (DMSO) or sham GSI-washout (GSI) were assessed using Cell Titer-Blue in quadruplicate independent cultures. (C) Formation of a 3D epidermal layer by SC2 cells in organotypic cultures in the presence of GSI or following GSI washout (WO). Cut sections of formalin-fixed, paraffin-embedded rafts were stained with H & E (hematoxylin and eosin) or with antibodies specific for ICN1 (activated intracellular NOTCH1) or the cell-cycle marker Ki67.

**Figure 2, Supplemental Figure 1.**
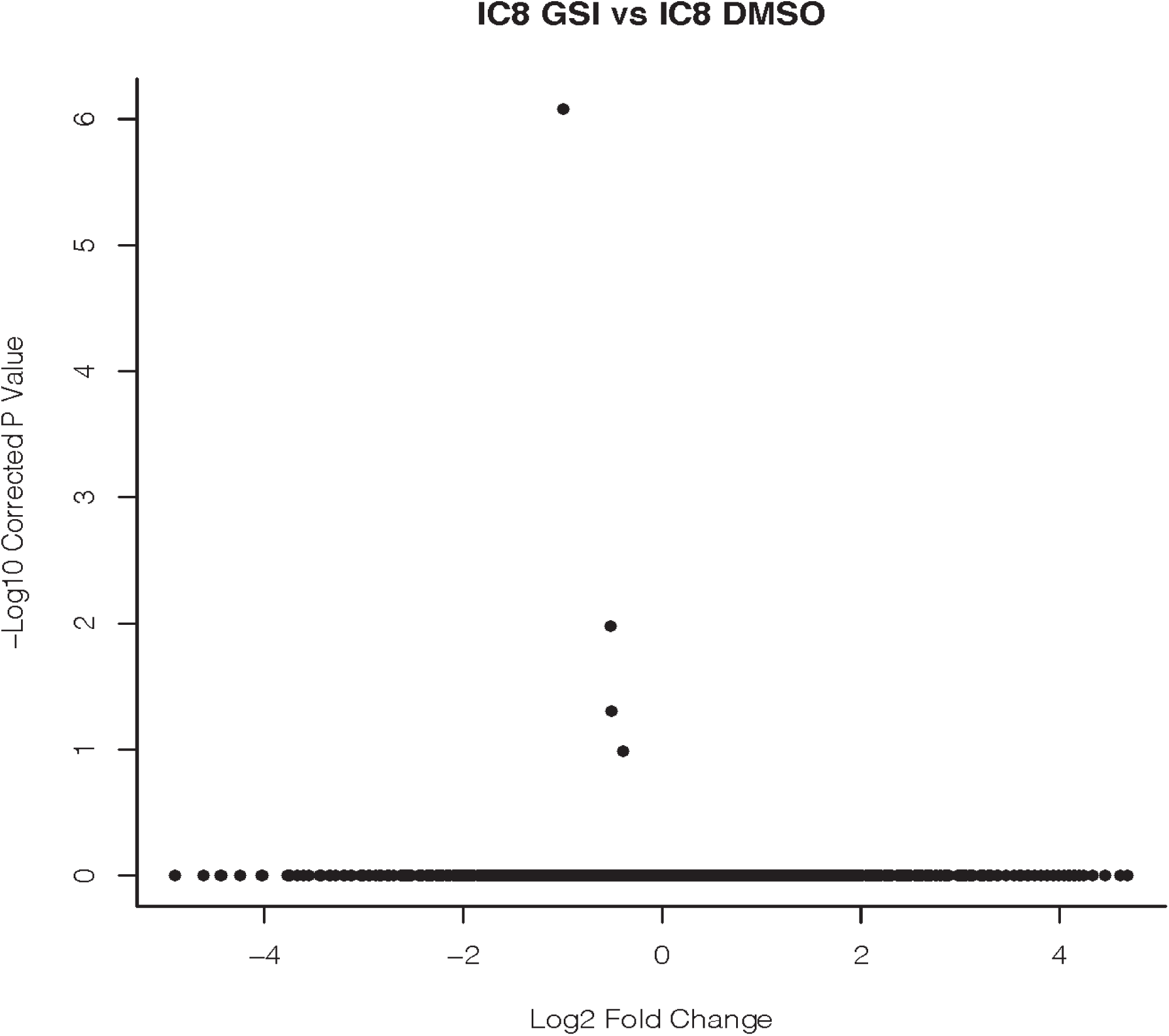
GSI has little effect on gene expression in IC8 cells. Duplicate cultures of IC8 cells were treated with GSI or vehicle (DMSO) for 24h and transcript abundance was assessed by RNA-seq. A volcano plot shows no differentially expressed genes using cutoffs of adjusted p-value < 0.0001 and log_2_ fold change > 1.

**Figure 4, Supplemental Figure 1.**
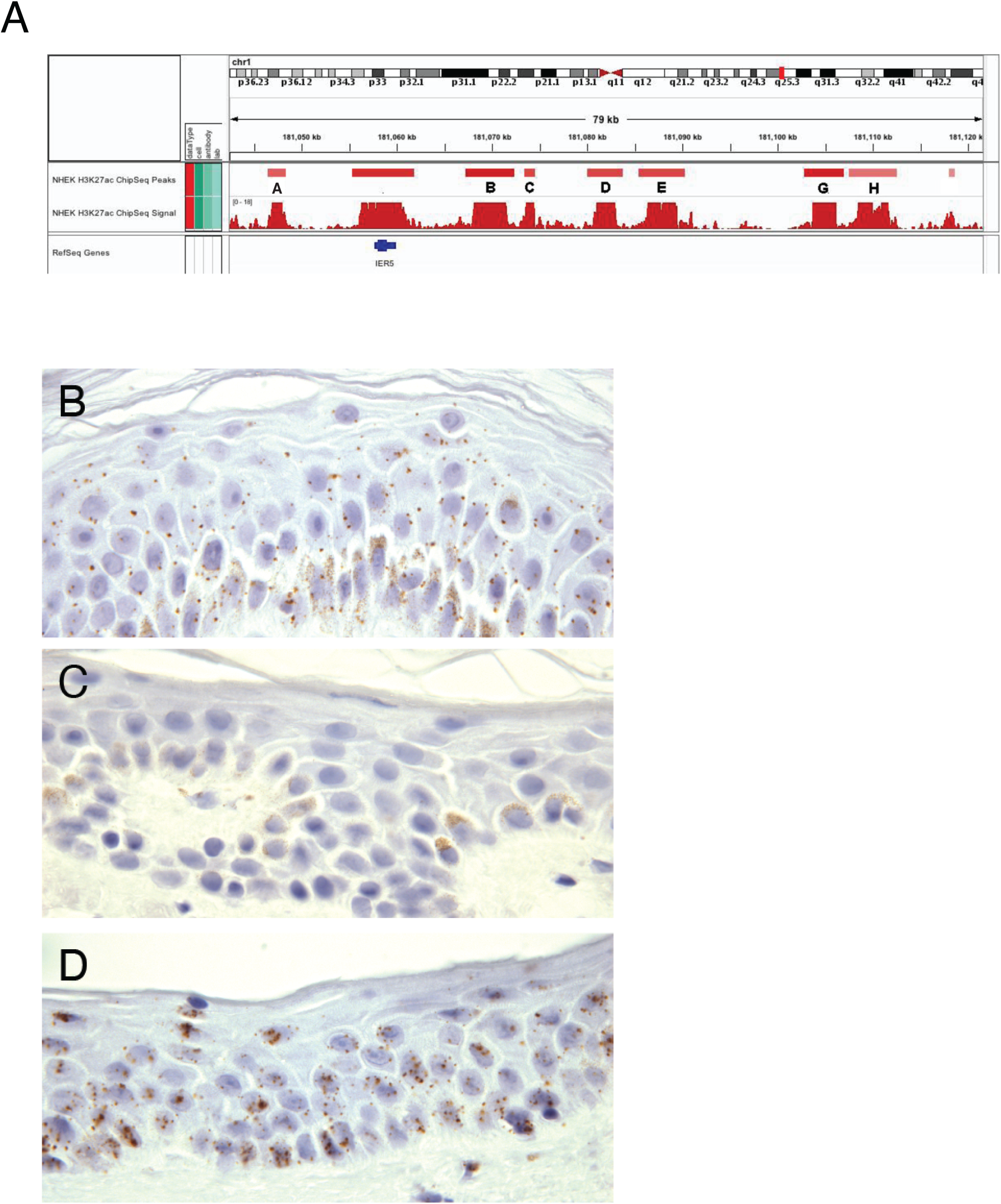
*IER5* enhancers and *IER5* expression in non-transformed keratinocytes. (A) H3K27ac landscapes near the *IER5* gene body in normal human epidermal keratinocytes (NHEK cells). ChIP-Seq data are from ENCODE. (B-D) Detection of *IER5* transcripts in normal human epidermis by in situ hybridization (ISH). B) *IER5*-specific probe; (C) *DapB*-specific negative control probe; (D) *PPIB*-specific positive control probe. Positive signals correspond to brown spots in cells that are counterstained with hematoxylin.

**Figure 5, Supplemental Figure 1.**
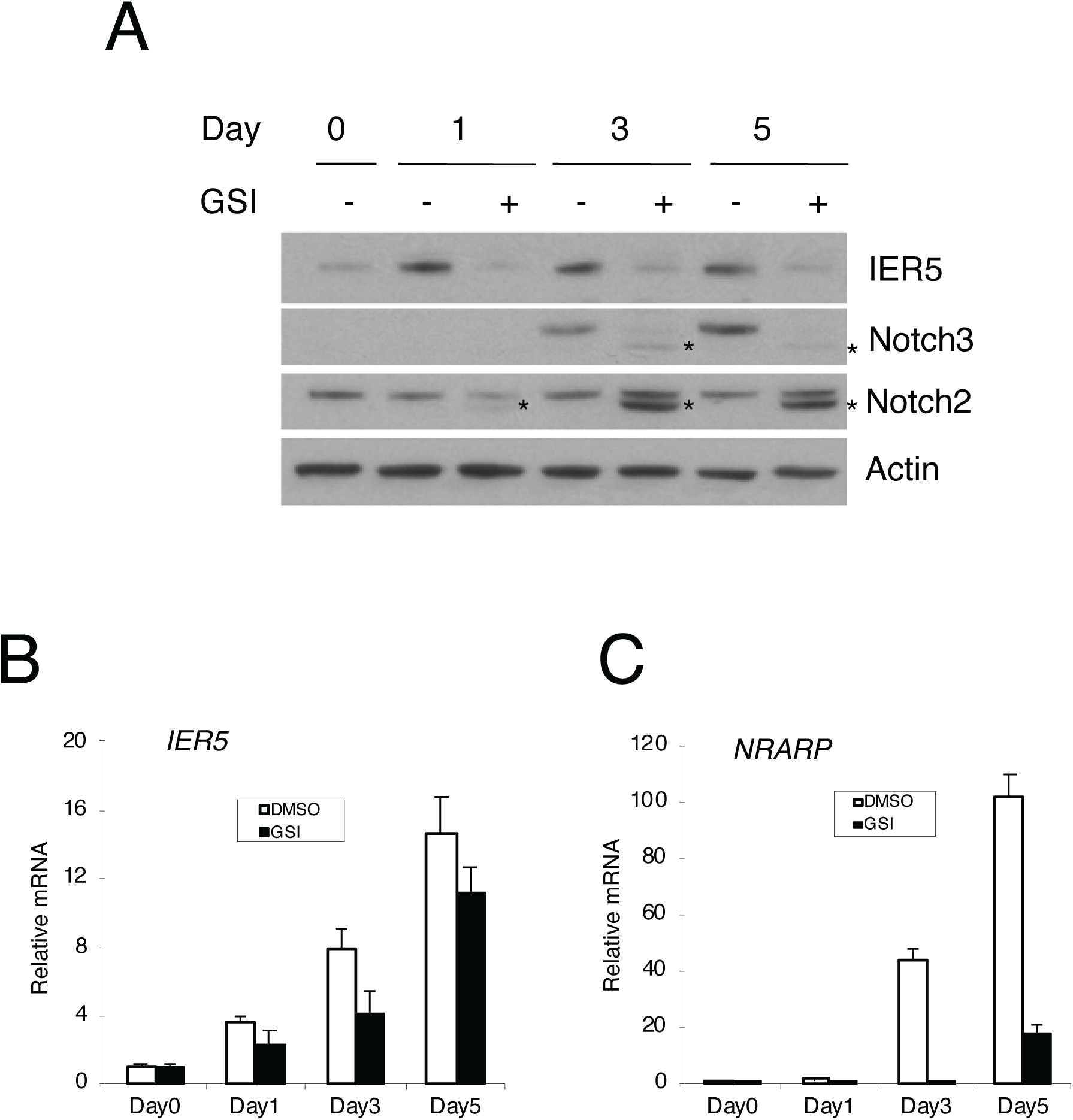
NOK1 cell differentiation is associated with Notch activation and increased *IER5* expression. (A) Western blot showing changes in IER5, NOTCH2, and NOTCH3 polypeptides following transfer to differentiation medium in the absence (-) and presence (+) of GSI. Increased levels of smaller NOTCH2 and NOTCH3 (denoted by asterisks) consistent with ADAM-metalloprotease cleaved products seen in differentiation medium in the presence of GSI are denoted with asterisks. (B) Changes in *IER5* and *NRARP* expression in the presence of vehicle (DMSO) and GSI) of Notch activation after transfer to differentiation medium. Transcript abundance was measured in experimental triplicates by RT-PCR and normalized against GAPDH. Error bars represent standard deviations of the mean.

## Acknowledgements and Author Contributions

### Funding

JCA is supported by the Harvard Ludwig Institute and the Michael A. Gimbrone Chair in Pathology at Brigham and Women’s Hospital and Harvard Medical School. SCB is supported by NIH award R35 CA220340. JMR is a fellow of the Leukemia and Lymphoma Society. JAM is supported by NIH award P01 CA203655.

### Author contributions

LP, WL, JMR, CJ, JM, TT, and GOA were responsible for acquisition, analysis, and interpretation of data. MEL was responsible for analysis and interpretation of data. APS, SCB, and JCA were responsible for experimental conception and design as well as interpretation of data. All authors contributed to drafting of the manuscript, have approved the content of the submission, and agree to be accountable for the work reported herein.

### Data and materials availability

All genome-wide data sets will be made available through GEO following publication. Cell lines, primer sequences, gRNA sequences, and other reagents/methodology will be made available on request.

## Supplemental Material

**Supplemental Table 1**. Differentially expressed genes after 4h of Notch activation, SC2 cells.

**Supplemental Table 2**. Differentially expressed genes after 24h of Notch activation, SC2 cells.

**Supplemental Table 3**. Differentially expressed genes after 72h of Notch activation, SC2 cells.

**Supplemental Table 4**. Unclustered GO annotations of Notch-sensitive genes, SC2 cells, at 72h of Notch activation.

**Supplemental Table 5**. Clustered GO annotations of Notch-sensitive genes, SC2 cells, at 72h of Notch activation.

**Supplemental Table 6**. Unclustered GO annotations of *IER5*-dependent Notch-sensitive genes, SC2 cells.

**Supplemental Table 7**. Clustered GO annotations of *IER5*-dependent Notch-sensitive genes, SC2 cells.

**Supplemental Table 8**. IER5-interacting proteins identified by tandem affinity purification.

